# Ionic Exposure History Shapes Inner Nuclear Membrane Voltage and Chromatin Texture Responses

**DOI:** 10.64898/2026.06.23.733978

**Authors:** Hamid Sediqi, Juanita Mathews, Giovanni de Nola, Abigail K. R. Lytton-Jean, Michael Levin

## Abstract

While bioelectricity is increasingly recognized as an important regulator of cell function and morphogenesis, the field has almost exclusively focused on plasma membrane states. Voltage across the inner nuclear membrane (INM) has been proposed as a potential regulator of nuclear function, but how it responds to extracellular ionic perturbations and whether it relates to chromatin organization remain unclear. Here, we targeted the ratiometric genetically encoded voltage indicator ASAP3-R3 to SUN2-associated nuclear membranes in intact NRK cells and combined INM voltage measurements with Gray-Level Co-Occurrence Matrix (GLCM)-based chromatin texture analysis. Reporter localization was confirmed by fluorescence imaging and electron microscopy, and functional validation in isolated nuclei showed that sodium-potassium pump inhibition produced INM depolarization consistent with Goldman-Hodgkin-Katz (GHK)-based prediction. We then used our validated construct to determine the response of V_nuc_ and chromatin texture to changing ionic conditions via two exposure methods, gradual (ramped) exposure or direct application. In intact cells, ramping different sets of ionic solutions of decreasing sodium/increasing potassium, decreasing sodium, increasing potassium, or decreasing chloride induced INM hyperpolarization and coordinated changes in chromatin texture, including increased contrast and entropy, reduced homogeneity, and reduced nuclear area. These effects were strongly path-dependent, with nuclear responses shaped by the history and order of ionic exposure: sodium and potassium responses emerged most clearly during ramping exposure, whereas reducing chloride by direct exposure showed a more pronounced response profile. Direct changes in sodium exposure produced limited electrical and chromatin-texture effects, while direct potassium exposure altered chromatin texture and nuclear area without significantly changing V_Nuc_. Importantly, shifting baseline chromatin state in either direction, through Trichostatin-A (TSA)-induced chromatin relaxation or sodium azide/2-deoxy-D-glucose-induced compaction, blunted ion-associated V_nuc_ and chromatin responses across sodium, potassium, and chloride conditions. Together, these findings identify the nucleus as a dynamic, ion-responsive electro-structural system in which INM voltage and chromatin organization are functionally coupled, and in which both ionic trajectory and pre-existing chromatin state shape the magnitude of the nuclear response.

## 1. Introduction

Recent decades have seen an explosion of interest in developmental and non-neuronal bioelectricity, revealing that ion fluxes and membrane voltage participate in information processing far beyond excitable tissues ^3–7^. Early work identified resting membrane potential (Vmem) as an important regulator of mitosis and neoplastic behavior, and subsequent studies have implicated V_mem_ in the control of proliferation, differentiation, migration, apoptosis, and gene regulation^3,5,8,9^. Bioelectric signals are now understood not only as physiological readouts, but also as instructive regulators of pattern formation, regeneration, and cancer-related cell-state transitions in vivo ^3–5,9^. Although most of this literature has focused on the plasma membrane, it has long been noticed that intracellular organelles also possess a bioelectrical dimension ^11^.

Especially intriguing was the early discovery of voltage across the nuclear envelope (NE), because it suggested that the nucleus itself might be electrically distinct from the surrounding cytoplasm. In a seminal study, Loewenstein and Kanno used microelectrodes in gland-cell nuclei of *Drosophila flavorepleta* and reported that the nucleus-cytoplasm boundary sustains both a measurable electrical resistance and a resting potential, with the nucleoplasm approximately 15 mV negative relative to the cytoplasm ^12^. Importantly, both the potential and resistance collapsed when the nuclear membrane was experimentally perforated, indicating that these electrical properties depended on the integrity of the nuclear boundary rather than representing an artifact of the recording geometry ^13,14^. These early observations established the basic point that the nucleus is not necessarily electrically continuous with the cytoplasm, and that the NE can behave as a selectively permeable boundary with its own electrical properties ^12,13^.

Over the following decades, electrophysiological studies began to reveal possible mechanisms for these early observations. Mazzanti and colleagues used patch-clamp recording on isolated mouse pronuclei and identified K^+^-selective channels in the NE with multiple conductance states, the largest of which was approximately 200 pS, and proposed that these channels contribute to nuclear membrane potential by regulating K^+^ movement across the NE ^14^. Additional studies then identified chloride-selective channels in isolated rat liver nuclei ^15^, distinct ionic conductance reconstituted from inner and outer nuclear membranes ^16^, and later functional ion-channel repertoires at the nuclear periphery more broadly ^11^. These findings were consolidated by later reviews into the idea of an “electrical dimension” of the NE, in which the nuclear boundary supports non-trivial ion selectivity and voltage behavior rather than acting as a purely passive extension of the endoplasmic reticulum (ER) ^11,13^.

In addition to passive ion channels, the NE can also establish ionic gradients through active transport. Garner showed that Na^+^/K^+^ gradients exist between the NE lumen and both the cytoplasm and nucleoplasm, and attributed these gradients to Na,K-ATPases located in the nuclear envelope, arguing that nuclear pores are not freely permeable to Na^+^ and K^+1^. Later work further linked nuclear ion transport to nuclear signaling by showing that Na^+^/Ca2^+^ exchanger activity at the nuclear envelope helps regulate nucleoplasmic Ca^2+^ homeostasis during differentiation ^17^. Together with broader reviews of nuclear channels and transporters, these studies support the view that the nuclear periphery actively contributes to nuclear ion homeostasis rather than passively compartmentalizing it ^1,11,13,17^.

At the same time, the idea of a nuclear membrane potential has always faced an obvious conceptual challenge: how can such a voltage exist given the large diameter of the nuclear pore complex (NPC)? At first glance, the NPC seems as though it should electrically short-circuit the nucleus by providing a wide aqueous conduit between nucleoplasm and cytoplasm. However, this objection assumes that the NPC behaves as an empty hole, which is not how the modern transport literature describes it. The NPC is instead a selective and dynamic permeability barrier filled with FG-repeat nucleoporins, whose transport properties are modulated by transport receptors and by the physiological state of the cell ^18–20^. Direct studies have also shown that pore permeability is responsive to ionic conditions, including Ca^2+^, while structural work has shown that NPC diameter can dilate and constrict in living cells depending on membrane mechanics and metabolic state ^21–23^. Thus, the presence of a large pore architecture does not by itself eliminate the possibility of nuclear voltage; rather, it means that any nuclear electrical state must be understood in the context of a regulated, finite-conductance transport barrier rather than a simple open short ^11,13,18–23^.

A related point is that nuclear voltage does not need to be conceptualized solely as one scalar difference between a perfectly mixed nucleoplasm and the cytoplasm. The NE creates three coupled routes for ion movement: across the outer nuclear membrane, across the inner nuclear membrane, and through the NPC. Accordingly, local ionic gradients and electrical states can in principle exist across one membrane relative to the perinuclear space even while some nucleocytoplasmic exchange still occurs through nuclear pores ^11,13,14^. This is especially relevant for the inner nuclear membrane (INM), which is the membrane most directly linked to the nuclear lamina, peripheral chromatin, and genome organization ^24,25^. In this sense, the key issue is not whether pores are present, but whether conductance across the nuclear boundary is selective, finite, and regulated enough to allow electrical states to emerge and persist. The electrophysiological and transport literature indicates that it is ^1,11,13,14,18–23^.

An additional point is that the nuclear interior is not a simple, well-stirred aqueous compartment in which ions and macromolecules equilibrate instantaneously. Early live-cell studies showed that chromatin undergoes constrained diffusional motion within limited nuclear subregions rather than exploring the entire nucleus freely, and that this mobility is further restricted by association with structures such as the nucleolus and nuclear periphery ^26,27^. Work from Elizabeth Hinde and colleagues extended this view by showing that molecular flow through the nucleus is strongly shaped by chromatin architecture. Using pair-correlation analysis, they demonstrated DNA-dependent molecular flow and showed that chromatin density regulates the diffusive path and temporal dynamics of an inert protein within the nucleus ^28,29^. More recent mechanical measurements add a complementary perspective by showing that chromatin behaves as a viscoelastic gel-like structure, whereas the nucleoplasm behaves as a softer viscoelastic phase rather than a freely mixed liquid ^30^. These observations do not prove that the NPC preserves a voltage gradient by itself, but they do argue against an overly simplified view in which the pore opens into a perfectly mixed sink. Instead, they support the idea that ion movement and electrical equilibration within the nucleus occur in a crowded, structured, and diffusion-constrained environment ^26–30^.

This picture is further supported by the growing literature on nuclear phase separation. Many nuclear processes, including transcriptional regulation, DNA replication, and RNA processing, occur within biomolecular condensates that locally enrich proteins and RNAs at defined genomic sites rather than allowing uniform mixing throughout the nucleus ^31^. Phase separation does not by itself explain how a nuclear membrane potential is generated or maintained, but it strengthens the broader argument that the nuclear interior is physically heterogeneous and locally compartmentalized. In that context, finite pore conductance together with constrained intranuclear diffusion, chromatin architecture, and condensate-based compartmentalization makes the idea of local or sustained electrical states at the nuclear envelope more plausible than would be expected from pore diameter alone ^18–23,26–31^.

The electrophysiological question is especially important because the NE is also a major organizer of genome architecture. The INM is compositionally distinct from the ONM, is closely associated with nuclear lamina, and contributes to chromatin organization and gene regulation ^24,25,32^. Chromatin itself is also highly sensitive to ionic conditions. Nucleosome arrays compact in response to defined salt environments, sodium and potassium do not influence higher-order chromatin folding identically, and nucleosomes retain a substantial negative electrostatic field even after DNA wraps around histones ^33–35^. These observations raise the possibility that ionic perturbations capable of altering voltage at the nuclear envelope may also influence chromatin architecture, and conversely, that baseline chromatin state may shape how strongly the nucleus responds electrically to ionic change ^24,25,33–35^.

Despite this conceptual overlap, direct interrogation of voltage specifically at the INM in intact mammalian cells remains limited. Much of the nuclear electrophysiology literature has focused on isolated nuclei, channel identification, calcium handling, or general NE transport properties ^1,11,13–17^. What has remained less clear is which extracellular ionic perturbations are sufficient to change INM voltage in intact cells, and whether such electrical changes are accompanied by coordinated changes in chromatin organization. To address the latter question, a quantitative measure of chromatin architecture is needed alongside the voltage readout. Gray-Level Co-Occurrence Matrix (GLCM) analysis provides such an approach. Originally developed by Haralick and colleagues, GLCM is a second-order texture method that quantifies how often pairs of grayscale intensities occur at a defined spatial offset ^36^. Rather than measuring only average fluorescence intensity, it captures how intensity values are arranged relative to one another across the image. This is particularly useful for chromatin because two nuclei with similar mean DNA staining can nonetheless differ markedly in the spatial organization of dense and sparse chromatin regions. Features derived from the co-occurrence matrix, such as contrast, entropy, and homogeneity, therefore offer a quantitative way to assess nuclear texture and detect changes in chromatin organization that may not be apparent by visual inspection alone ^36–38^.

A second technical requirement is a voltage reporter that can be localized to the nuclear membrane itself. Kim and colleagues recently developed ASAP3-R3, a ratiometric genetically encoded voltage indicator in which the voltage-sensitive ASAP3 reporter is fused to the voltage-insensitive red fluorophore mCyRFP3, allowing the green/red ratio to report voltage while correcting for motion-related artifacts ^39^. In parallel, earlier work in plants showed that ArcLight derivatives can be targeted to nuclear membranes using SUN2-based localization strategies ^40^, and mammalian studies established SUN2 as an inner nuclear membrane protein ^41^. These prior advances made it feasible to adapt a ratiometric genetically encoded voltage indicator (GEVI) for INM voltage measurements in mammalian cells.

Here, we targeted ASAP3-R3 to the INM using a SUN2 truncated sequence and linker, then used this reporter to examine, for the first time, how defined extracellular ionic perturbations affect INM voltage in intact normal rate kidney fibroblast (NRK-49F) cells. In parallel, we quantified chromatin texture using GLCM analysis of Hoechst-labelled nuclei. Our results show that increasing extracellular potassium or decreasing extracellular chloride induce INM hyperpolarization and coordinated changes in chromatin texture, whereas sodium perturbation produces bidirectional responses that depend on exposure history. Direct exposure to lower extracellular sodium levels causes depolarization and a more homogeneous chromatin texture, while gradual exposure (ramping) to lower sodium induces hyperpolarization and a compaction-like chromatin signature. We further show that potassium responses are strongly order-dependent, with ramped exposure producing effects not observed under direct exposure to the same endpoint condition, revealing a kind of non-Markovian memory effect dynamic. Finally, experimentally relaxing or compacting chromatin blunts the normal voltage and chromatin responses to ionic challenge, indicating that baseline chromatin state constrains nuclear electrical responsiveness. Together, these findings support the view that the INM is a regulated bioelectric membrane with path dependence on a time-scale of 10 minutes, and that nuclear voltage dynamics are closely coupled to chromatin organization.

## 2. Methods

### 2.1 Molecular Biology

The CAG ASAP3-R3 construct was subcloned from pc3.1-CAGGS-ASAP3-mCyRFP3 using NheI and HindIII and was a gift from Michael Lin (Addgene plasmid # 193323; http://n2t.net/addgene:193323; RRID:Addgene_193323). The fragment was cloned into a pENTR1A plasmid with a CAG promoter and multiple cloning site (MCS) followed by an SV40 poly(A) using the same sites as were used in excising the fragment from the parent plasmid. The NLS composed of amino acids 26 - 339 of the natively expressed inner nuclear membrane protein Sun2 was linked with a glycine serine linker to connect the NLS to the amino terminus of ASAP3-R3. In preparation for Gateway cloning, the NLS DNA sequence was first synthesized using Telesis BioXP and subsequently cloned into the pENTR ASAP3-R3 vector using Gibson Assembly (NEB E2611). The resulting pENTR1A CAG Sun2 ASAP3-R3 was then Gateway LR clonased (11791020, ThermoFisher) into the hyperactive piggyBac transposase-based, helper-independent, and self-inactivating delivery system, pmhyGENIE-3, a gift from Stefan Moisyadi ^42,43^. The pmhyGENIE-3 had a neomycin resistance gene in the backbone and the resulting plasmid HypG3 NeoBB Sun2 ASAP3-R3, was purified using a ZymoPURE Plasmid Miniprep Kit (D4210, Zymo Research) and used for subsequent transfections.

### 2.2 Cell culture

The normal rat kidney fibroblast cell line NRK-49F (CRL-1570, ATCC) was grown in Fluorobrite growth media (A1896701, Thermo Fisher) supplemented with 10% Fetal Bovine Serum (FBS) (16140071, Thermo Fisher), penicillin/streptomycin, and Glutamax (35050061, Thermo Fisher). Cells were maintained in a 37°C, 5% CO_2_ humidified incubator then passaged at 80% confluence by washing twice with with DPBS lacking calcium and magnesium (D837, Sigma-Aldrich), then dissociated with Accutase (25-058-CI, Corning)). Cells were cryopreserved using complete media with 10% dimethyl sulfoxide (DMSO) (D2438, Sigma). After thawing, cells were grown for at least a week before being used for experiments.

### 2.3 Transfection

NRK-49F cells were cultured in a 24-well plate at a confluence of 30% before being transfected with 500 ng of DNA plus P3000 and 1 uL of Lipofectamine 3000 (L3000008, Thermo Fisher) in Optimem (31985062, Thermo Fisher) according to manufacturer’s protocol. After 24 hours the cells were given fresh media, and after 48 hours the transfected cells were selected with Geneticin (1000 μg/ml). After 14 days cells were serially diluted to obtain clonally pure lines with acceptable expression and proliferation profiles.

### 2.4 Nuclei Isolation

Cells were dissociated from culture plates as described above, transferred to 15 mL Falcon tubes, and centrifuged at 0.2 rcf for 3 minutes. The medium was aspirated, and the cell pellet was resuspended in ice-cold DPBS before being centrifuged again at 0.2 rcf for 3 minutes. After removal of the DPBS, the tube was placed on ice, and cells were resuspended in 1 mL of ice-cold hypotonic buffer containing 10 mM Tris-Cl, pH 7.4, 10 mM NaCl, and 0.5 mM MgCl. Cells were incubated on ice in the dark for 60 minutes, then gently passed through a 27-gauge insulin syringe 5–10 times, taking care to avoid bubble formation. The suspension was examined under a microscope using a 40× objective after every fifth pass to monitor lysis and prevent over-disruption. Once approximately 70–90% of nuclei were released from the cytoplasm, the suspension was passed twice through a 40 µm cell strainer, using a fresh strainer for the second pass. The isolated nuclei were then resuspended in nuclear suspension buffer containing 145 mM KCl, 10 mM NaCl, 3 mM MgCl, and 20 mM Tris-HCl, pH 7.4, supplemented with 1 µM Hoechst 33342. Nuclei were subsequently divided into two groups: a control group treated with an equivalent volume of DMSO, and a treatment group exposed to 50 µM ouabain (Sigma-Aldrich, Cat. O3125-5G) and 250 µM strophanthidin (MedChemExpress, Cat. HY-114252).

### 2.5 V_mem_ Modulating Treatment

Cryopreserved Sun2 ASAP3-R3 stably transfected NRK-49F clones were thawed at least a week prior to experiments and cultured using complete Fluorobrite growth media. The cells were then cultured at 20 000 cells per well into a 96-well plate (Cellvis, Cat: P96-1.5P) 24 hours prior to imaging. Cells were then stained with Hoechst 33342 (1 µM) in complete Fluorobrite media for 10 minutes before being exposed to the osmotically balanced Na^+^/K^+^, Na^+^, K^+^, Cl-series of ion solutions as described by Bonzanni *et al*.^44^ and then imaged.

### 2.6 Imaging

Nuclear V_mem_ characterization was performed by acquiring widefield images using the ImageXpress HT-AI confocal microscope equipped with a 40x water objective. Hoechst-stained Nuclei were excited at 405 nm and emission was collected using the DAPI filter. Cells expressing ASAP3-R3-INM were excited at 488 nm and 586 nm, the green emission was collected using the FITC filter, while the red emission was collected using the Texas Red filter, respectively.

### 2.7 Global Chromatin Modulating Treatment

Global chromatin relaxation was induced by inhibiting histone deacetylase activity with Trichostatin A (TSA). For the sodium and potassium series, NRK-49F cells cultured in Cellvis 96-well plates were treated with 50 nM TSA for 24 h before imaging. For the chloride series, TSA was used at 50-200 nM. Control cells were exposed to medium containing an equivalent volume of DMSO for 24 h.

Global chromatin condensation was induced by ATP depletion using sodium azide and 2-deoxy-D-glucose (2-DG). NRK cells cultured in Cellvis 96-well plates were first incubated for 20 min in incomplete glucose-free DMEM (Thermo Fisher Scientific, Cat. A1443001) containing 10 mM sodium azide, 50 mM 2-DG, and 1 µM Hoechst 33342. Cells were then exposed for a further 10 min to glucose-free ionic solutions containing the same concentrations of sodium azide and 2-DG before imaging. Control cells were incubated for 20 min in complete FluoroBrite medium containing 1 µM Hoechst 33342 and were subsequently exposed to glucose-containing ionic solutions.

### 2.8 Image quality control and processing

Hoechst-stained nuclear images were used to generate ROI masks by applying a Gaussian blur (sigma=1), then thresholding using the RenyiEntropy algorithm in ImageJ. These images were subjected to quality control before texture analysis and Sun2 ASAP3-R3 fluorescence quantification to exclude blurred images. Blur detection was performed using a custom Python script implementing a Tenengrad-like focus metric. For each Hoechst image, the Sobel gradient magnitude was computed, and the mean gradient intensity was measured within each segmented nuclear ROI. ROIs touching the image border were excluded. Images containing fewer than five usable ROIs were also excluded from further analysis. An image-level focus score was then derived from the distribution of ROI focus scores using the 25th percentile, providing a conservative estimate of image sharpness. After all eligible images were scored, a global cutoff was defined as the 10th percentile of image-level focus scores across the dataset. Images with focus scores at or below this threshold were classified as blurry and excluded from subsequent analyses, whereas images above this threshold were retained.

### 2.9 V_mem_ Quantification

ROI masks of non-blurred image sets were then used to quantify Sun2 ASAP3-R3 Green and Red intensities. These values were saved in an csv file, and the Green/Red ratio of ASAP3-R3-INM was computed. To compute ΔV_nuc,_ the ΔG/R (%) was divided by 41% which is the magnitude of change per 100 mV, then multiplied by 100 ((ΔG/R)/41%*100). The data from the excel sheet were transferred to ‘Prism Graph Pad’ for statistical analysis and data visualization.

### 2.10 GLCM texture analysis

Texture analysis was performed on the retained Hoechst images using a custom Python pipeline implementing masked gray-level co-occurrence matrix (GLCM) analysis. Before feature extraction, each image was normalized to 16-bit intensity values using the 99th percentile of pixel intensities within the full nuclear mask. Specifically, the 99th percentile intensity of all mask-positive pixels in a given image was used as the normalization reference, and the image was rescaled such that this value corresponded to 65,535. Values exceeding this range were clipped. This normalization reduced variation in absolute signal intensity across images while preserving intranuclear intensity relationships relevant to chromatin texture.

To standardize the region used for texture analysis, a centroid-centered square patch was extracted from each ROI. Before final feature extraction, all ROIs were scanned to determine feasible patch sizes. For each ROI, the maximum even-valued box size that could fit entirely within the ROI was estimated using a chessboard distance transform. From this distribution, the largest box sizes compatible with 80%, 90%, and 100% of ROIs were determined, and a final even-valued box size was selected for analysis. Only ROIs in which the full square patch fit completely within the ROI mask were retained.

For each included ROI, texture was quantified from the extracted square patch using a masked GLCM approach. Pixel intensities were quantized into 8 gray levels before GLCM calculation. Co-occurrence matrices were computed at pixel distances of 2, 4, 8, and 12 pixels and across four angular directions (0°, 45°, 90°, and 135°). Only pixel pairs for which both pixels lay within the ROI mask were included. The resulting GLCMs were symmetrized and normalized to generate probability matrices. From each GLCM, three texture features were calculated: contrast, homogeneity, and entropy. Contrast was defined as

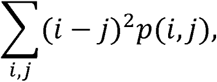

reflecting local gray-level variation. Homogeneity was defined as

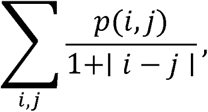

reflecting the extent to which co-occurrence probabilities were concentrated near the diagonal. Entropy was defined as

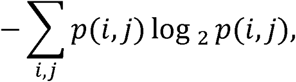

with a small constant added during computation to avoid numerical instability at zero-probability values. Texture features were first calculated at the individual ROI level. ROI-level values were then summarized per image by calculating the mean and standard deviation for each feature, together with the number of ROIs analyzed. These image-level values were subsequently aggregated to generate summary statistics for each experimental group. Separate summaries were generated for each GLCM distance. The change in contrast, entropy, and homogeneity calculated by deducted values from the first solution from the subsequent solutions.

## 3. Results

### 3.1 Targeting and Validation of INM-localized GEVI: ASAP3-R3

To characterize the voltage potential of the inner nuclear membrane (INM), we generated a nuclear-envelope-targeted ASAP3-R3 construct by fusing the SUN2 nuclear-targeting sequence (amino acids 26-339) to the N-terminus of ASAP3-R3 through a flexible glycine-serine linker (Figure 1A-i). Fluorescence imaging showed enrichment of the construct at the nuclear membrane, consistent with successful targeting of ASAP3-R3 to the nuclear periphery (Figure 1A-ii). Transmission electron microscopy further confirmed localization of ASAP3-R3-Nuc in the nuclear envelope, including within type I and type II nucleoplasmic reticulum invaginations extending from the nuclear envelope into the nucleoplasm (Figure 1B). These observations establish that the reporter is positioned at a biologically relevant nuclear membrane compartment suitable for monitoring INM voltage dynamics in intact cells.

**Figure 1.**
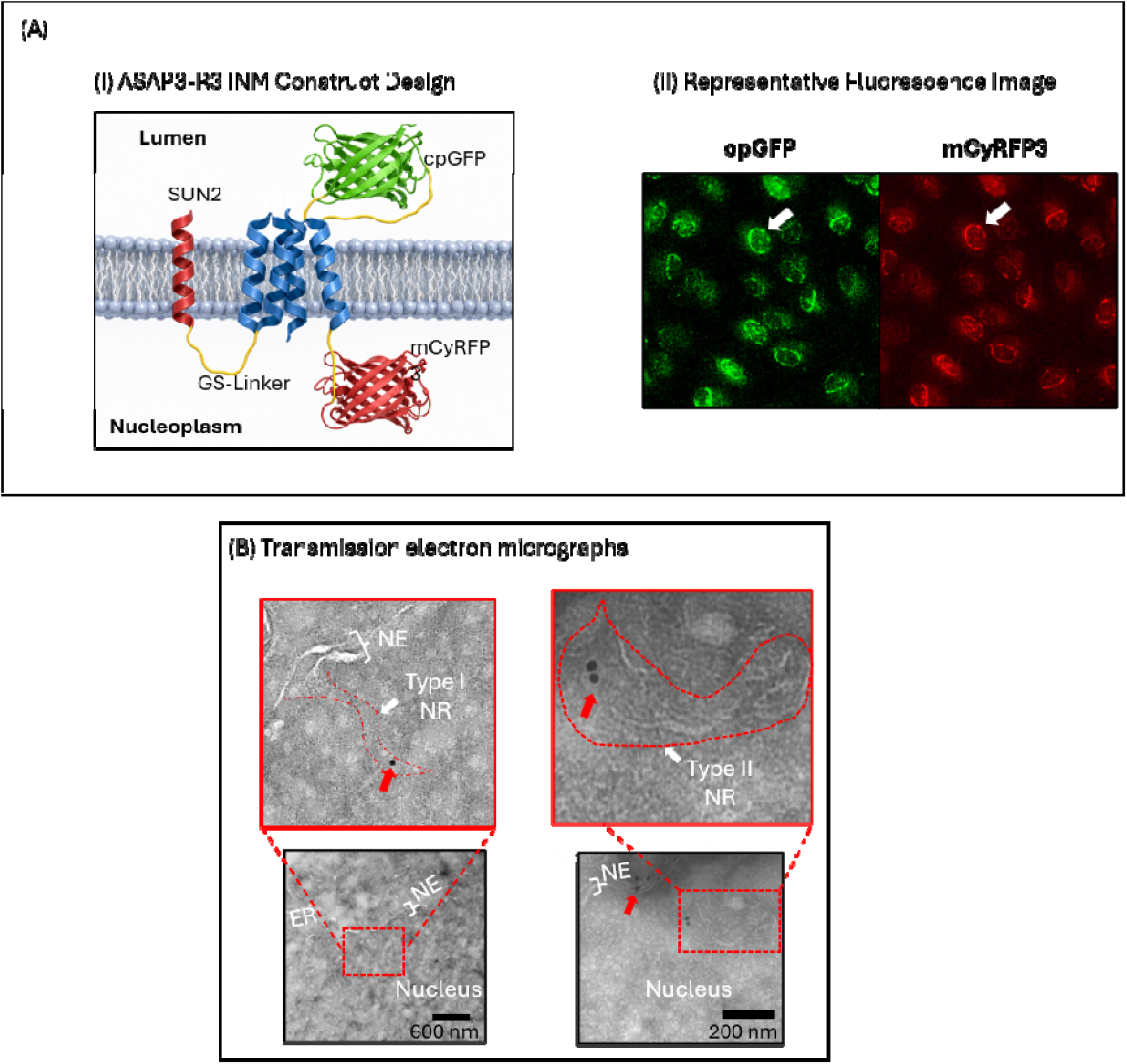
Successful localization of ASAP3-R3 to the inner nuclear envelope. (A-i) Schematic of the ASAP3-R3-Nuc construct design. The nuclear-targeting region of SUN2, comprising amino acids 26-339, was fused to the N-terminus of ASAP3-R3 through a flexible glycine-serine linker to preserve the fluorescent properties of cpGFP and mCyRFP3. (A-ii) Representative fluorescence micrographs showing cpGFP fluorescence in green and mCyRFP3 fluorescence in red. White arrows indicate enrichment of the construct at the nuclear membrane. (B) Electron micrograph showing localization of ASAP3-R3-Nuc within type I and type II nucleoplasmic reticulum invaginations of the nuclear envelope.

We next validated whether INM-ASAP3-R3 could detect a known perturbation of nuclear membrane potential. In isolated nuclei, inhibition of the INM sodium-potassium pump with strophanthidin and ouabain produced an approximately ^+^5 mV depolarization relative to DMSO-treated controls (p=0.003, n=10, one-sample test; Figure 2A). This experimentally measured shift was consistent with the estimated membrane potential calculated using the Goldman-Hodgkin-Katz (GHK) equation and reported nuclear sodium ^1^, potassium ^1^, and chloride concentrations ^2^, assuming standard permeability ratios reported for the plasma membrane: PK:PNa:PCl ≈ 1:0.03:0.5 ^10^ (Figure 2B). These findings support the use of INM-ASAP3-R3 as a reporter of INM voltage changes.

**Figure 2.**
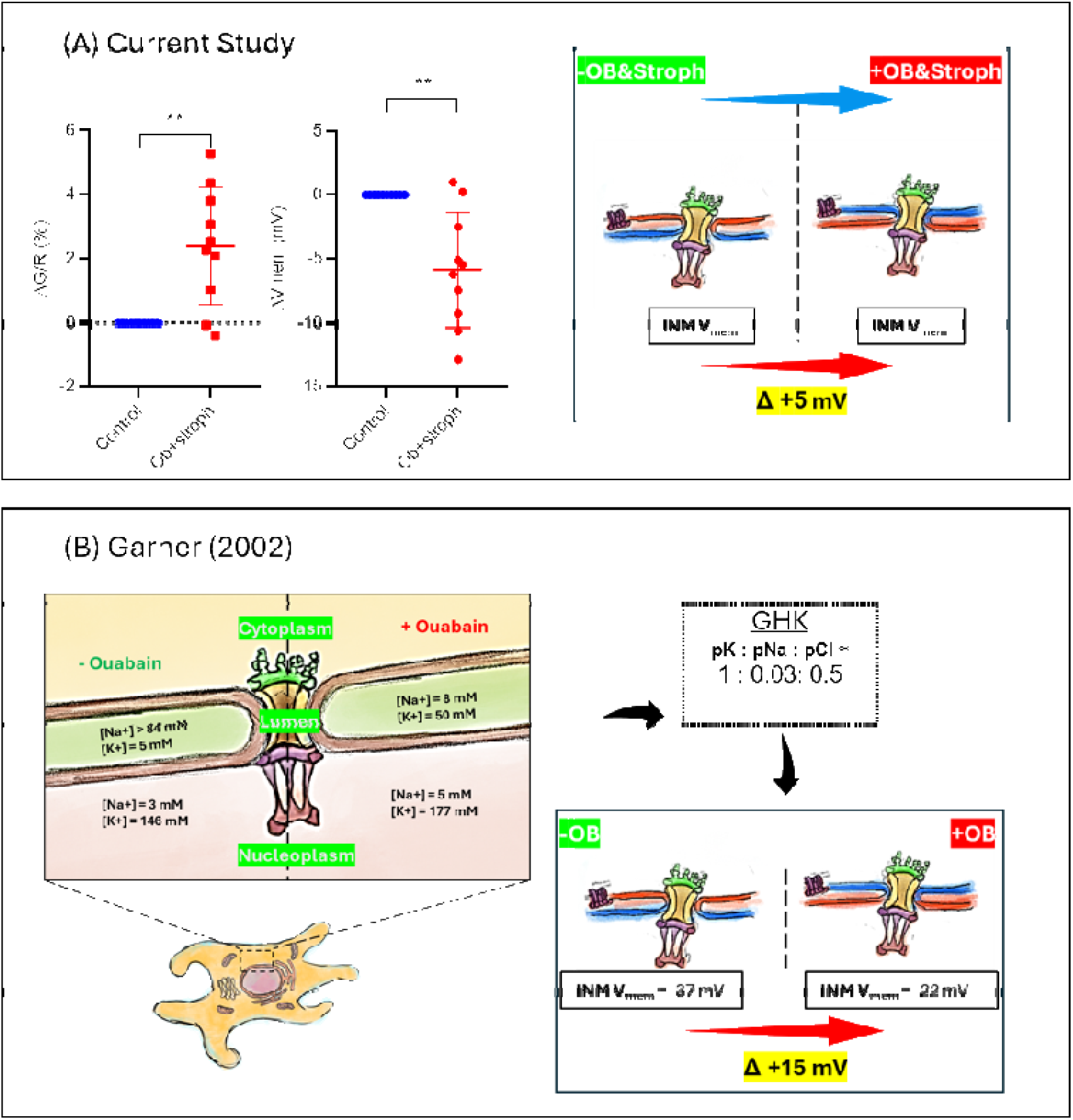
Validation of inner nuclear membrane ASAP3-R3. Isolated nuclei from NRK cells were loaded with 1 µM Hoechst 33342 and incubated at on ice for 15 min. Then, the nuclei were split into two groups - control were given equal volume of DMSO while the second group was given 250 nM strophanthidin and 50 µM ouabain. Nuclei were then dispensed into 96-well (Celvis) and left to incubate for 1 hour at room temperature to inhibit inner nuclear membrane sodium pump before imaging with the ImageXpress Confocal microscope. Hoechst 33342 was excited with 405 nm and emission was collected with DAPI filter, cpGFP of INM-ASAP3R3 was excited with 488 nm, and emission was collected with FITC filter, and RFP domain was excited at 586 nm and emissions were collected using Texas Red filter. Masks of isolated nuclei were generated using a filter for size and roundness. These masks were dilated to include nuclear envelope for quantification of Green/Red ratio. (A) Inhibition of the INM sodium-potassium pump resulted in ^+^5 mV depolarization (i) when normalized to DMSO-control nuclei. ((n=10, p =0.003, one-sample t-test, error bars represent standard deviation). (B) This value aligns with estimated V_mem_from the GHK equation using sodium and potassium concentrations reported by Garner ^1^ and chloride concentration reported by Satish ^2^, assuming standard permeabilities pK:pNa:pCl ≈ 1: 0.03: 0.5 ^10^.

### 3.2 Mixed Sodium/Potassium Ramping Uncovers Stateful Coupling of INM Hyperpolarization and Chromatin Texture Remodeling

We next asked whether extracellular ionic perturbations alter INM voltage and whether these changes are accompanied by measurable shifts in chromatin organization. We exposed the cells to the different ion solutions using two methods; a gradual (ramped) exposure that incrementally changed the ion solutions until the final end point concentration was reached, or a direct exposure that immediately switched out the external solution for the final end point concentration. In a mixed sodium/potassium calibration series, direct exposure to a single sodium/potassium condition did not significantly change nuclear membrane potential (Vnuc) relative to baseline (p>0.05, n=10, one-way ANOVA; Figure 3A-i). Consistent with this, GLCM analysis showed no significant changes in contrast, entropy, or homogeneity under the direct-exposure paradigm (p>0.05, n=10, one-sample test, FDR-corrected; Figure 3B-i). However, nuclear area increased significantly following direct exposure (0.0004<p<0.02, n=10, one-sample test, FDR-corrected; Figure 3B-i). In contrast, sequential ramping exposure to solutions containing increasing potassium and decreasing sodium concentrations produced significant graded hyperpolarization of the INM, reaching approximately −9 mV at 45 mM sodium/100 mM potassium (p<0.005, n=10, one-way ANOVA; Figure 3A-ii). This electrical response was accompanied by significant graded increases in chromatin contrast and entropy, together with a graded decrease in homogeneity (0.01<p<0.05, n=10, one-sample test, FDR-corrected; Figure 3B-ii). Nuclear area analysis showed a significant graded decrease across the ramping series (0.0001<p<0.05, n=9, one-sample test, FDR-corrected; Figure 3B-ii). Thus, the mixed calibration series revealed that INM voltage responses are not triggered simply by endpoint ionic conditions but emerge more clearly during progressive ionic transitions and are accompanied by coordinated changes in chromatin texture.

**Figure 3.**
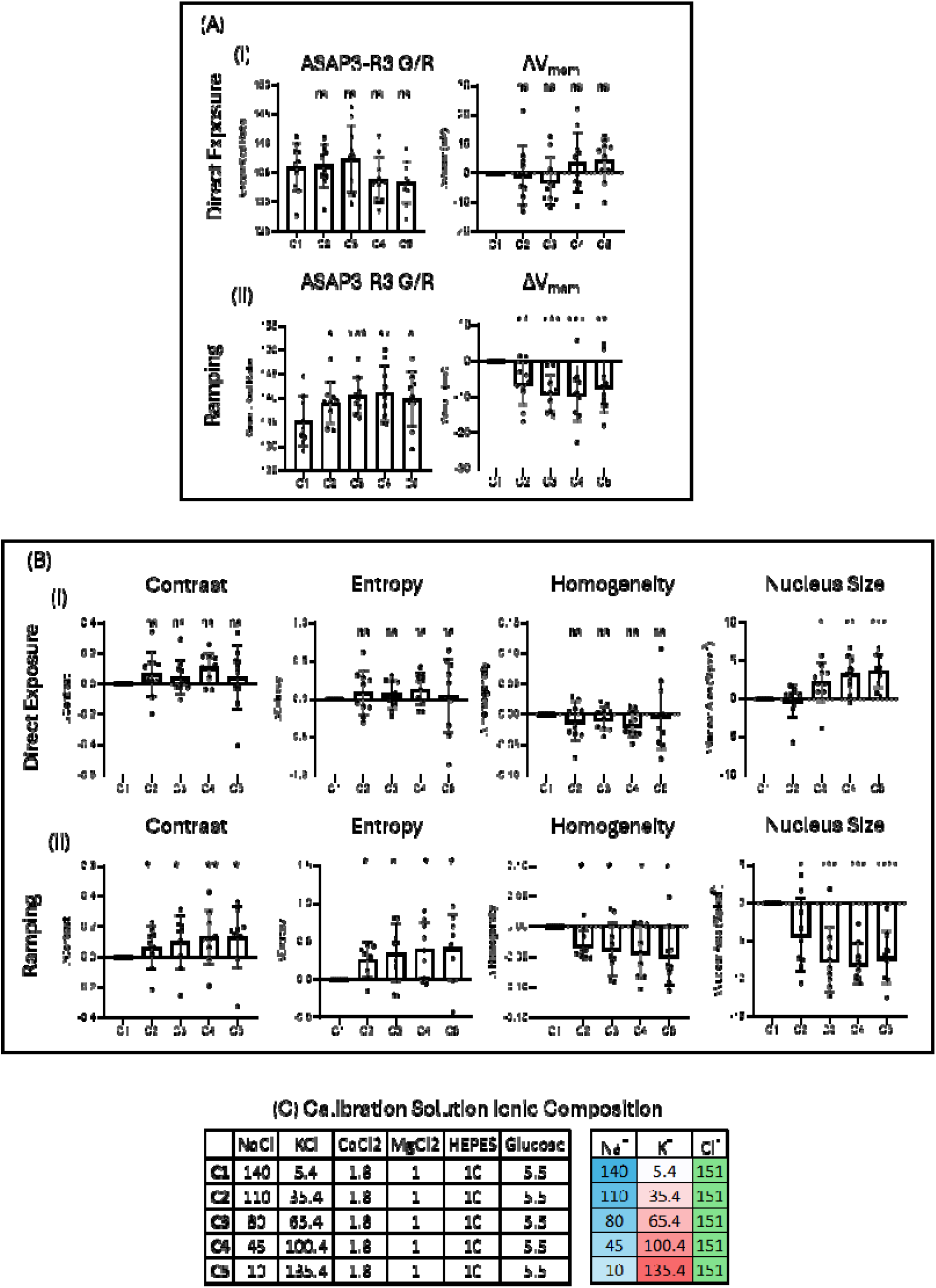
Calibration Series Causes INM Hyperpolarization and Chromatin Compaction. Intact NRK cells were cultured in Cellvis 96-well plates for 24 h before imaging at 37 °C and 5% CO_₂_. Cells were stained with Hoechst 33342 (1 µM) in complete FluoroBrite medium, after which the medium was replaced with mixed ionic solutions. In direct-exposure experiments, cells were exposed to a single sodium/potassium condition, whereas in ramping experiments cells were sequentially exposed to solutions containing increasing potassium and decreasing sodium concentrations. (A) ASAP3-R3 measurements at the inner nuclear membrane (INM) showed no significant change in nuclear membrane potential (Vnuc) in (i) direct-exposure experiments (p>0.05, n=10, one-way ANOVA), but revealed significant graded hyperpolarization in (ii) ramp-up experiments, reaching approximately-9 mV at 45mM sodium/100 mM potassium (p<0.005, n=10, one-way ANOVA). (B-i) GLCM analysis showed no significant changes in contrast, entropy, or homogeneity following direct exposure (p>0.05, n=10, one-sample Wilcoxon test, FDR-corrected), however showed significant nuclear swelling (0.02 <p>0.0004, n=10, one-sample test, FDR-corrected). (B-ii) In ramp-up experiments, GLCM analysis revealed a significant graded increase in contrast and entropy, accompanied by a graded decrease in homogeneity (0.05<p<0.01, n=10, one-sample Wilcoxon test, FDR-corrected) and nuclear area (0.05<p<0.0001, n=9, one-sample test, FDR-corrected). (C) Ionic composition of calibration solution. Together, these findings are consistent with increased chromatin compaction, greater architectural complexity, and reduced chromatin homogeneity. Error bars represent standard deviation.

### 3.3 Ramping Sodium Reduction Couples INM Hyperpolarization and Chromatin Texture Remodeling

To determine whether INM voltage responses to sodium reduction depended on the temporal order of ionic exposure, we next compared direct and ramping sodium-only perturbations. Under direct-exposure conditions, acute presentation of sodium-modified solutions produced only limited changes in Vnuc. Most sodium conditions did not significantly alter VNucrelative to baseline, although a significant response was detected at 28 mM sodium (p<0.05, n=14, one-way ANOVA; Figure 4A-i). Consistent with this limited voltage response, direct sodium exposure did not significantly alter GLCM-derived chromatin contrast, entropy, or homogeneity (p>0.05, n=9, one-sample test, FDR-corrected; Figure 4B-i).

**Figure 4.**
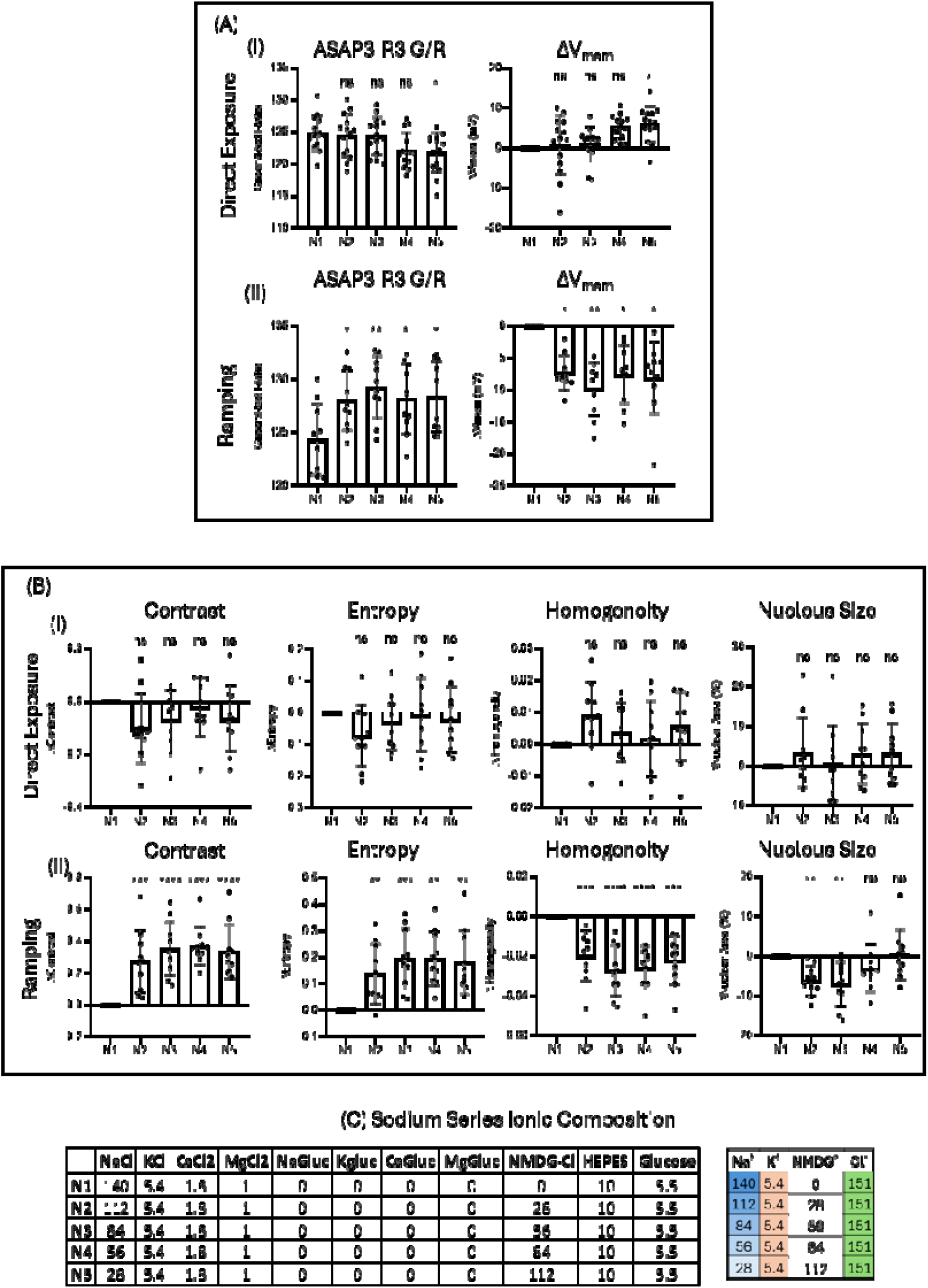
Sodium series induces INM hyperpolarization and chromatin compaction. Intact NRK cells were cultured in Cellvis 96-well plates for 24 h before imaging at 37 °C and 5% CO_₂_. Cells were stained with Hoechst 33342 (1 µM) in complete FluoroBrite medium, after which the medium was replaced with potassium-containing solutions. In direct-exposure experiments, cells were exposed to a single potassium concentration, whereas in ramping experiments cells were sequentially exposed to solutions containing progressively lower sodium concentrations. (A) ASAP3-R3 measurements at the inner nuclear membrane (INM) revealed no significant change in nuclear membrane potential (Vnuc) in (i) direct-exposure experiments for all solutions except for N5 (p<0.05, n=14, one-way ANOVA). Conversely, a significant graded hyperpolarization was observed in (ii) ramping experiments (p<0.05, n=10, one-way ANOVA), reaching approximately-10 mV at 84 mM sodium. (B-i) GLCM analysis revealed no significant changes in contrast, entropy, or homogeneity following direct exposure (p>0.05, n=9, one-sample test, FDR-corrected). (B-ii) In ramp-up experiments, GLCM analysis showed a significant graded increase in contrast and entropy, accompanied by a graded decrease in homogeneity (p<0.05, n=9, one-sample Wilcoxon test, FDR-corrected) and nuclear are at 84 mM sodium. (C) Ionic composition of potassium series. Together, these findings are consistent with increased chromatin compaction, greater architectural complexity, and reduced chromatin homogeneity. Error bars represent standard deviation.

In contrast, when nuclei experienced sodium reduction as a progressive ramp, lowering sodium concentration produced significant graded hyperpolarization of the INM, reaching approximately −10 mV at 84 mM sodium (p<0.05, n=10, one-way ANOVA; Figure 4A-ii). This ramping-dependent voltage response was accompanied by coordinated chromatin texture changes, including graded increases in contrast and entropy and a graded decrease in homogeneity (p<0.05, n=9, one-sample test, FDR-corrected; Figure 4B-ii). Nuclear area was analyzed separately and showed a significant reduction at 84 mM sodium.

Thus, sodium reduction did not produce a fixed INM response determined solely by the sodium concentration present at the time of measurement. Instead, the effect of sodium depended on the temporal context in which the nucleus encountered the ionic condition. Acute sodium reduction produced only limited electrical and chromatin-texture effects, whereas progressive sodium reduction revealed robust INM hyperpolarization accompanied by chromatin compaction-associated texture changes. These findings suggest that sodium-associated INM voltage dynamics are history-dependent, with the nuclear response shaped by prior ionic exposure rather than by instantaneous sodium concentration alone.

### 3.4 Ramping Potassium Elevation Reveals History-Dependent Coupling Between INM Voltage and Chromatin Texture Remodeling

To determine whether INM voltage responses depended only on the current ionic condition or instead were influenced by the prior ionic history of the nucleus, we compared direct and ramping exposure to potassium-only solutions. Under direct exposure, presentation higher potassium conditions did not significantly alter Vnuc relative to baseline (p>0.05, n=11, one-way ANOVA; Figure 5A-i). However, the same direct potassium exposure altered chromatin texture, decreasing contrast and entropy while increasing homogeneity (p<0.05, n=11, one-sample test, FDR-corrected; Figure 5B-i). Nuclear area also increased significantly following direct potassium exposure (p<0.01, n=11, one-sample test, FDR-corrected; Figure 5B-i).

**Figure 5.**
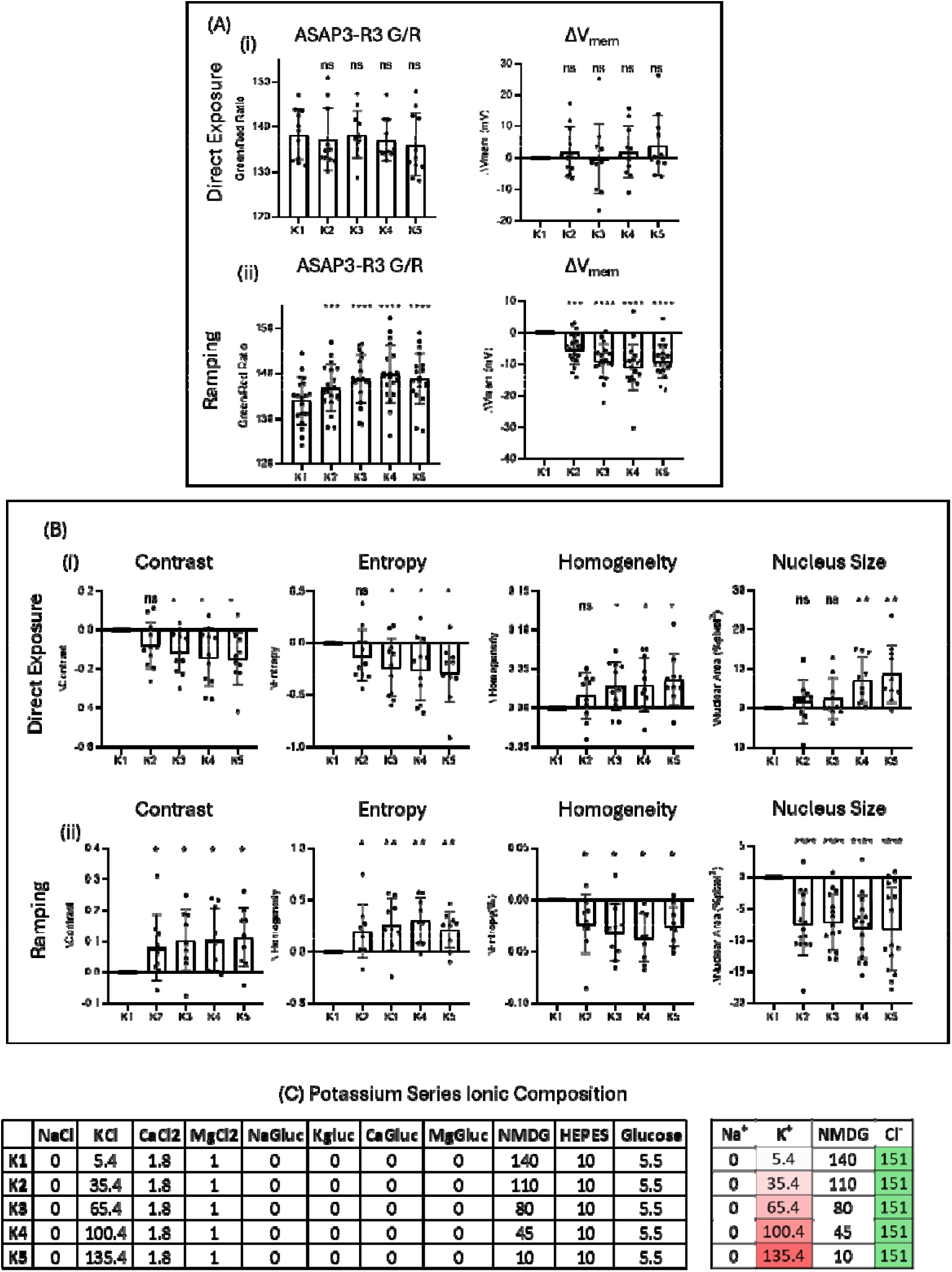
Potassium series induces INM hyperpolarization and chromatin compaction. Intact NRK cells were cultured in Cellvis 96-well plates for 24 h at 37 °C and 5% CO_₂_ before imaging. Cells were stained with Hoechst 33342 (1 µM) in complete FluoroBrite medium, after which the medium was replaced with potassium-containing solutions. In direct-exposure experiments, cells were exposed to a single potassium concentration, whereas in ramping experiments cells were sequentially exposed to solutions containing progressively higher potassium concentrations. (A) ASAP3-R3 measurements at the inner nuclear membrane (INM) revealed no significant change in nuclear membrane potential (Vnuc) in (i) direct-exposure experiments (p>0.05, n=11, one-way ANOVA) but showed significant graded hyperpolarization in (ii) ramping experiments (p<0.0001, n=19, one-way), reaching approximately-10 mV at 100 mM potassium. (B-i) GLCM analysis revealed significant decrease in contrast, entropy, and increase in homogeneity and nuclear area following direct exposure (p>0.05, n=11, one-sample Wilcoxon test, FDR-corrected). (B-ii) In ramping experiments, GLCM analysis showed a significant graded increase in contrast and entropy, accompanied by a graded decrease in homogeneity (p<0.05, n=9, one-sample test, FDR-corrected) and nuclear area (p<0.0001, n=19, one-sample test, FDR-corrected). (C) Ionic composition of potassium series. Together, these findings are consistent with increased chromatin compaction, greater architectural complexity, and reduced chromatin homogeneity. Error bars represent standard deviation.

In contrast, when nuclei encountered the potassium condition as part of a progressive ramp, increasing extracellular potassium produced significant graded INM hyperpolarization, reaching approximately −10 mV at 100 mM potassium (p<0.0001, n=19, one-way ANOVA; Figure 5A-ii). This voltage response was accompanied by graded increases in chromatin contrast and entropy, a graded decrease in homogeneity (p<0.05, n=9, one-sample test, FDR-corrected; Figure 5B-ii), and a significant graded decrease in nuclear area (p<0.0001, n=19, one-sample test, FDR-corrected; Figure 5B-ii).

Thus, the final potassium condition did not produce a fixed INM voltage response. Instead, the effect of potassium depended on the temporal order in which the nucleus experienced the ionic environment. Acute exposure to potassium altered chromatin texture without significantly changing Vnuc, whereas progressive potassium ramping produced robust INM hyperpolarization together with coordinated chromatin compaction-associated texture changes. These findings suggest that INM voltage dynamics are history-dependent and stateful, rather than simply reflecting the instantaneous potassium concentration.

### 3.5 Chloride-Dependent INM Voltage-Chromatin Coupling Diverges from Sodium and Potassium Response Patterns

To determine whether chloride-associated INM voltage responses were also shaped by their history of prior ionic exposure, we next compared direct and ramping chloride-only perturbations. Unlike the sodium and potassium series, direct reduction of chloride concentration produced significant graded INM hyperpolarization (0.001<p<0.05, n=10, one-way ANOVA; Figure 6A-i). This voltage response was accompanied by coordinated chromatin texture changes, including significant graded increases in contrast and entropy together with a graded decrease in homogeneity (0.01<p<0.05, n=10, one-sample test, FDR-corrected; Figure 6B-i). Nuclear area also decreased significantly with decreasing chloride concentration (0.001<p<0.05; Figure 6B-i).

**Figure 6.**
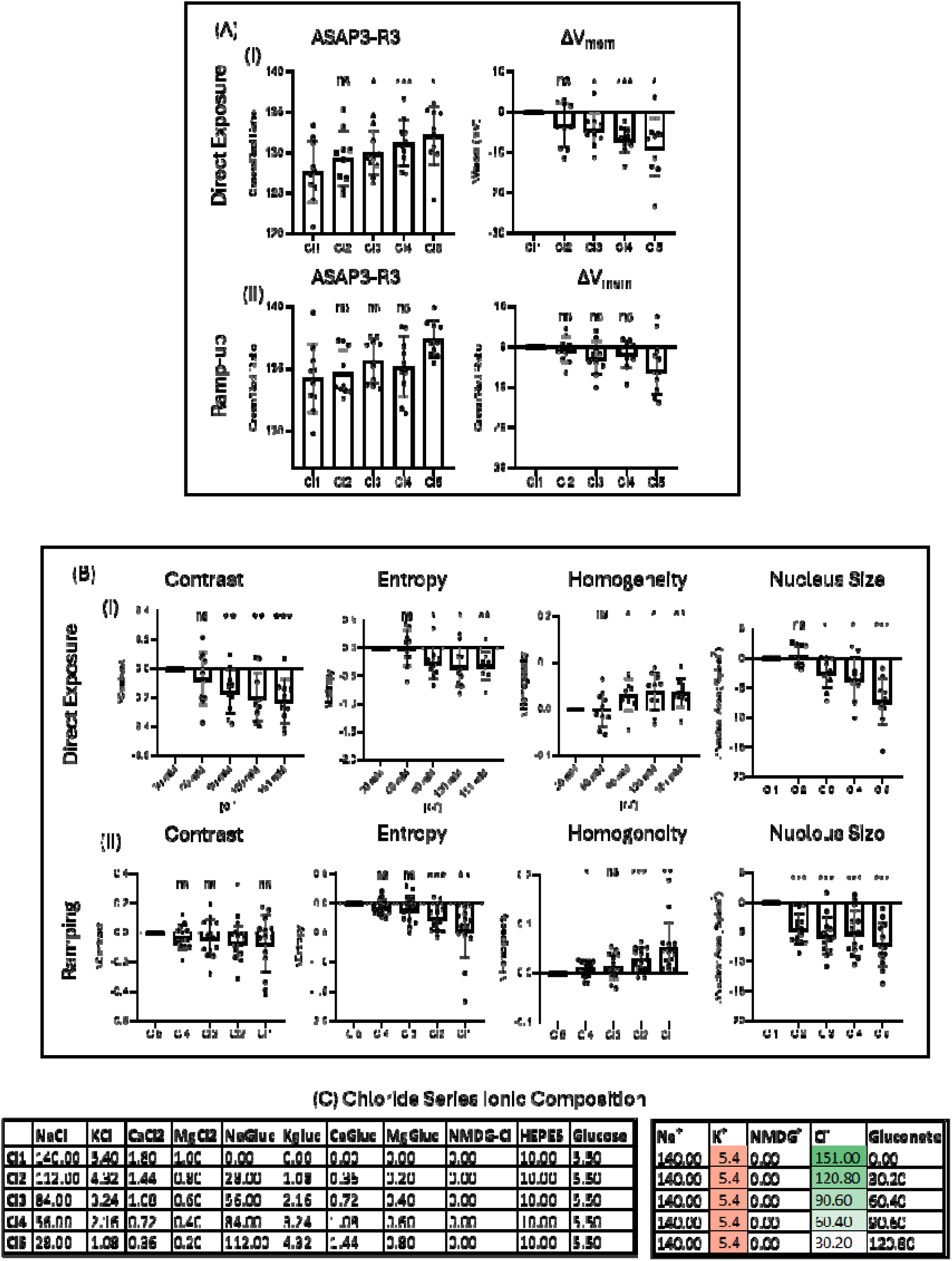
Chloride series induces INM hyperpolarization and chromatin compaction. Intact NRK cells were cultured in Cellvis 96-well plates for 24 h at 37 °C and 5% CO_₂_ before imaging. Cells were stained with Hoechst 33342 (1 µM) in complete FluoroBrite medium, after which the medium was replaced with chloride-modified solutions. In direct-exposure experiments, cells were exposed to a single chloride concentration, whereas in ramping experiments cells were sequentially exposed to solutions containing progressively lower chloride concentrations. (A) ASAP3-R3 measurements at the inner nuclear membrane (INM) showed significant graded hyperpolarization in response to decreasing chloride concentration in (i) direct-exposure experiments (0.05<p<0.001, n=10, one-way ANOVA). (ii) Ramping experiments revealed only 30 mM Chloride solution resulted in hyperpolarization of INM (p<0.05, n=10, one-way ANOVA), reaching approximately-6 mV. (B-i) GLCM analysis of direct-exposure experiments revealed significant graded increases in contrast and entropy, together with a graded decrease in homogeneity (0.01<p<0.05, n=10, one-sample Wilcoxon test, FDR-corrected). (B-ii) In ramping experiments, GLCM analysis showed no significant change in contrast, but did reveal a graded increase in entropy and a graded decrease in homogeneity (p<0.01, n=14, one-sample test, FDR-corrected) and nuclear area (p<0.001, n=10. (C) Ionic composition of chloride series. Together, these findings are consistent with increased chromatin compaction, greater architectural complexity, and reduced chromatin homogeneity.

In contrast, when nuclei experienced chloride reduction as a progressive ramp, the voltage response was more restricted. Only the 30 mM chloride condition produced significant INM hyperpolarization, reaching approximately −6 mV (p<0.05, n=10, one-way ANOVA; Figure 6A-ii). Despite this more limited Vnuc response, ramping chloride exposure still altered chromatin texture, with entropy increasing and homogeneity decreasing in a graded manner, while contrast did not change significantly (p<0.01, n=14, one-sample test, FDR-corrected; Figure 6B-ii). Nuclear area also decreased significantly during the chloride ramp (p<0.001, n=10, one-sample test, FDR-corrected; Figure 6B-ii).

Thus, chloride reduction did not produce the same exposure-history pattern observed for sodium and potassium. Whereas sodium and potassium responses were most clearly revealed during progressive ramping, chloride reduction produced a stronger graded voltage response during direct exposure and a more condition-specific response during ramping. These findings indicate that chloride-associated INM voltage and chromatin responses are also history-dependent, but follow a distinct exposure-history profile from sodium and potassium.

### 3.6 Chromatin Relaxation Blunts Sodium-Associated INM Voltage Responses

Because sodium, potassium, and chloride perturbations were each associated with coordinated changes in V_nuc_ and chromatin texture, we next asked whether the pre-existing chromatin state influences the magnitude of the nuclear electrical response. To test this, we first induced global chromatin relaxation with trichostatin A (TSA). During sodium ramping experiments, TSA treatment significantly reduced the V_nuc_ response to decreasing sodium concentration relative to DMSO-treated controls (p<0.001, n=10, two-way ANOVA; Figure 7A). GLCM analysis showed no significant difference in contrast between TSA-treated and DMSO-treated cells (Figure 7B-i), whereas entropy and homogeneity responses were significantly blunted in TSA-treated cells (p<0.0001, n=10, two-way ANOVA; Figure 7B-ii,iii). Nuclear area analysis also revealed significant blunting following TSA treatment (Figure 7B-iv). Thus, chromatin relaxation diminished sodium-associated V_nuc_ responses and selectively dampened the accompanying chromatin-texture dynamics.

**Figure 7.**
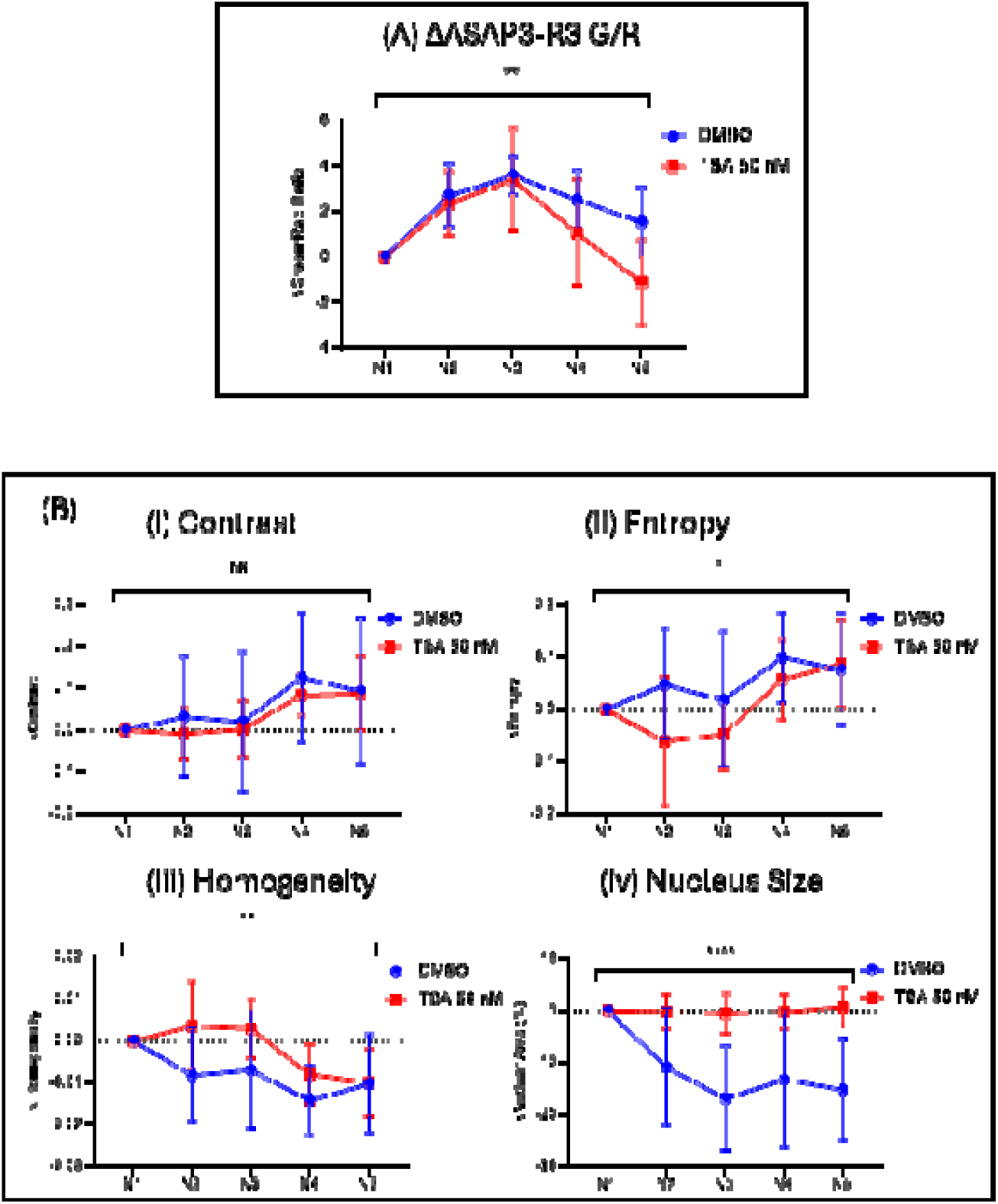
Global chromatin relaxation blunts sodium V_nuc_ responses and associated chromatin dynamics. Intact NRK cells were cultured in Cellvis 96-well plates for 48 h at 37 °C and 5% CO_₂_ before imaging. Cells were treated with TSA (50 nM) or an equivalent volume of DMSO for 24 h before imaging. Cells were then stained with Hoechst 33342 (1 µM) in complete FluoroBrite medium, and the medium was replaced with sodium-containing solutions. During ramping experiments, cells were sequentially exposed to solutions containing progressively lower sodium concentrations (A) ASAP3-R3 measurements at the inner nuclear membrane (INM) showed that TSA treatment significantly reduced the V_nuc_ response to decreasing sodium concentration relative to DMSO-treated controls (p<0.001, n=10, two-way ANOVA). (B) GLCM analysis similarly revealed no significant difference in (i) contrast, but (ii) entropy and (iii) homogeneity was significantly blunted in TSA-treated cells (p<0.0001, n=10, two-way ANOVA). Change in nuclear area was also significantly blunted in TSA treated cells. These findings are consistent with reduced chromatin compaction dynamics and diminished potassium-associated changes in chromatin organization. Error bars represent standard deviation.

### 3.7 Chromatin Compaction Blunts Sodium-Associated INM Voltage Responses

We next examined whether global chromatin compaction also altered sodium-associated responses. Sodium azide/2-DG treatment significantly reduced the V_nuc_ response to decreasing sodium concentrations relative to controls (p<0.001, n=10, two-way ANOVA; Figure 8A). This treatment also significantly blunted sodium-associated changes in contrast, entropy, and homogeneity (p<0.0001, n=10, two-way ANOVA; Figure 8B-i-iii). Changes in nuclear area also appeared blunted in sodium azide/2-DG-treated cells. These findings show that forcing chromatin toward a compacted state also reduces sodium-associated INM electrical responsiveness and chromatin reorganization.

**Figure 8.**
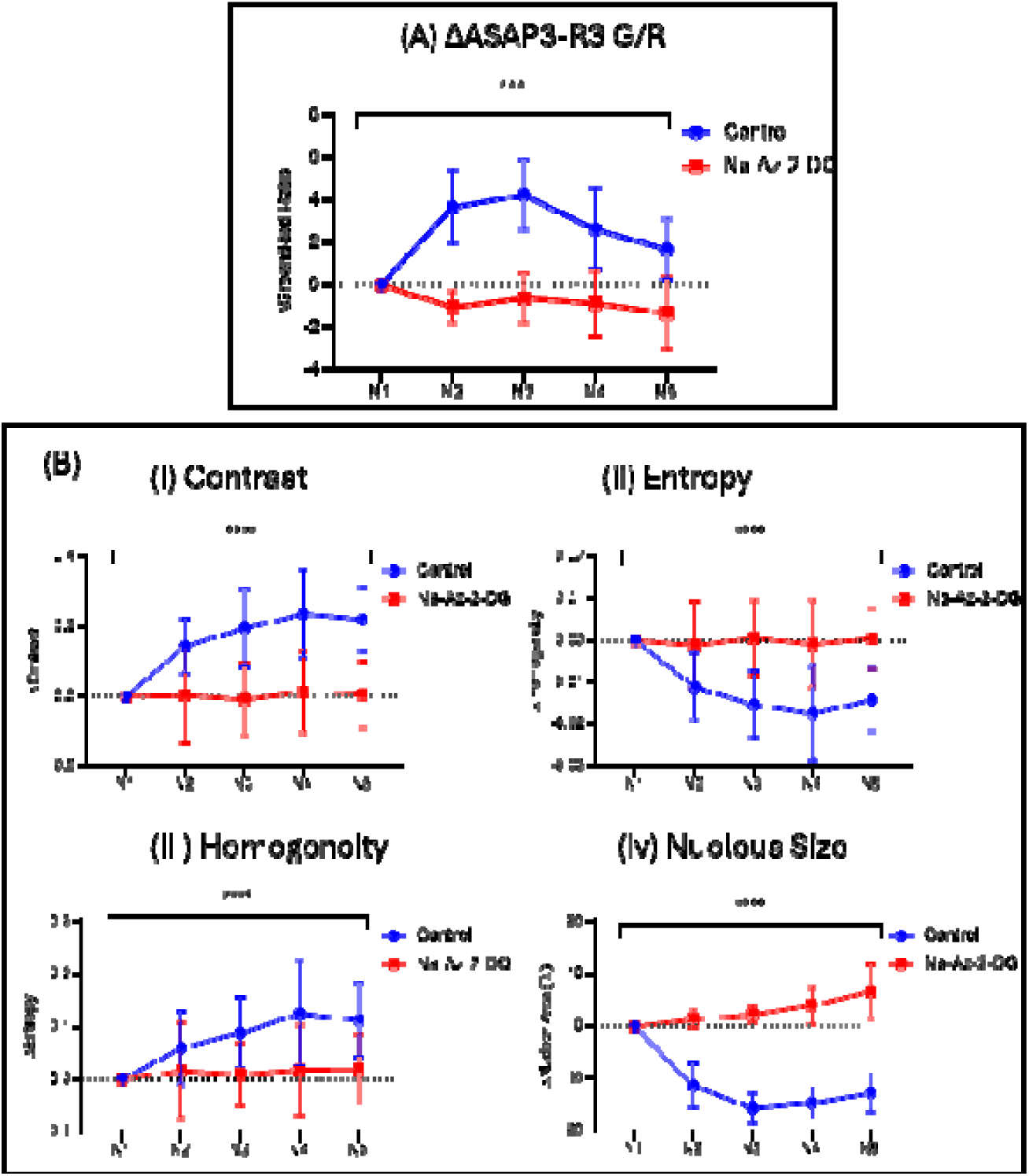
Global chromatin compaction blunts sodium-induced V_nuc_ responses and associated chro atin dynamics. Intact NRK cells were cultured in Cellvis 96-well plates for 24 h before imaging at 37 °C and 5% CO_₂_. Cells were incubated for 20 min in control complete Fluorobrite medium containing Hoechst 33342 (1 µM) alone or in incomplete DMEM media without glucose containing Hoechst 33342 (1 µM), sodium azide (10 mM), and 2-deoxy-D-glucose (2-DG; 50 mM). Cells were subsequently exposed to a ramping sodium series lacking glucose and containing sodium azide and 2-DG for an additional 10 min before imaging, whereas control cells were exposed to a glucose-containing ramping sodium series. (A) ASAP3-R3 measurements at the inner nuclear membrane (INM) showed that sodium azide/2-DG treatment significantly reduced the V_nuc_ response to decreasing sodium concentrations relative to controls (p<0.001, n=10, two-way ANOVA). (B) GLCM analysis similarly revealed significantly blunted responses in (i) contrast, (ii) entropy, and (iii) homogeneity in treated cells (p<0.0001, n=5, two-way ANOVA). Nuclear area also appeared blunted in Na-Az 2DG treated cells. These results are consistent with reduced sodium-associated changes in chromatin organization following induction of global chromatin compaction. Error bars represent standard deviation.

### 3.8 Chromatin Relaxation Blunts Potassium-Associated INM Voltage Responses

We then tested whether chromatin relaxation affected potassium-associated responses. Relative to DMSO-treated controls, TSA treatment significantly blunted the V_nuc_ response to increasing potassium concentration (p<0.0001, n=10, two-way ANOVA; Figure 9A). TSA treatment also significantly blunted potassium-associated changes in chromatin contrast, entropy, and homogeneity (p<0.0001, n=10, two-way ANOVA; Figure 9B-i-iii). Thus, chromatin relaxation diminished not only the structural response of chromatin itself, but also the accompanying electrical responsiveness of the INM.

**Figure 9.**
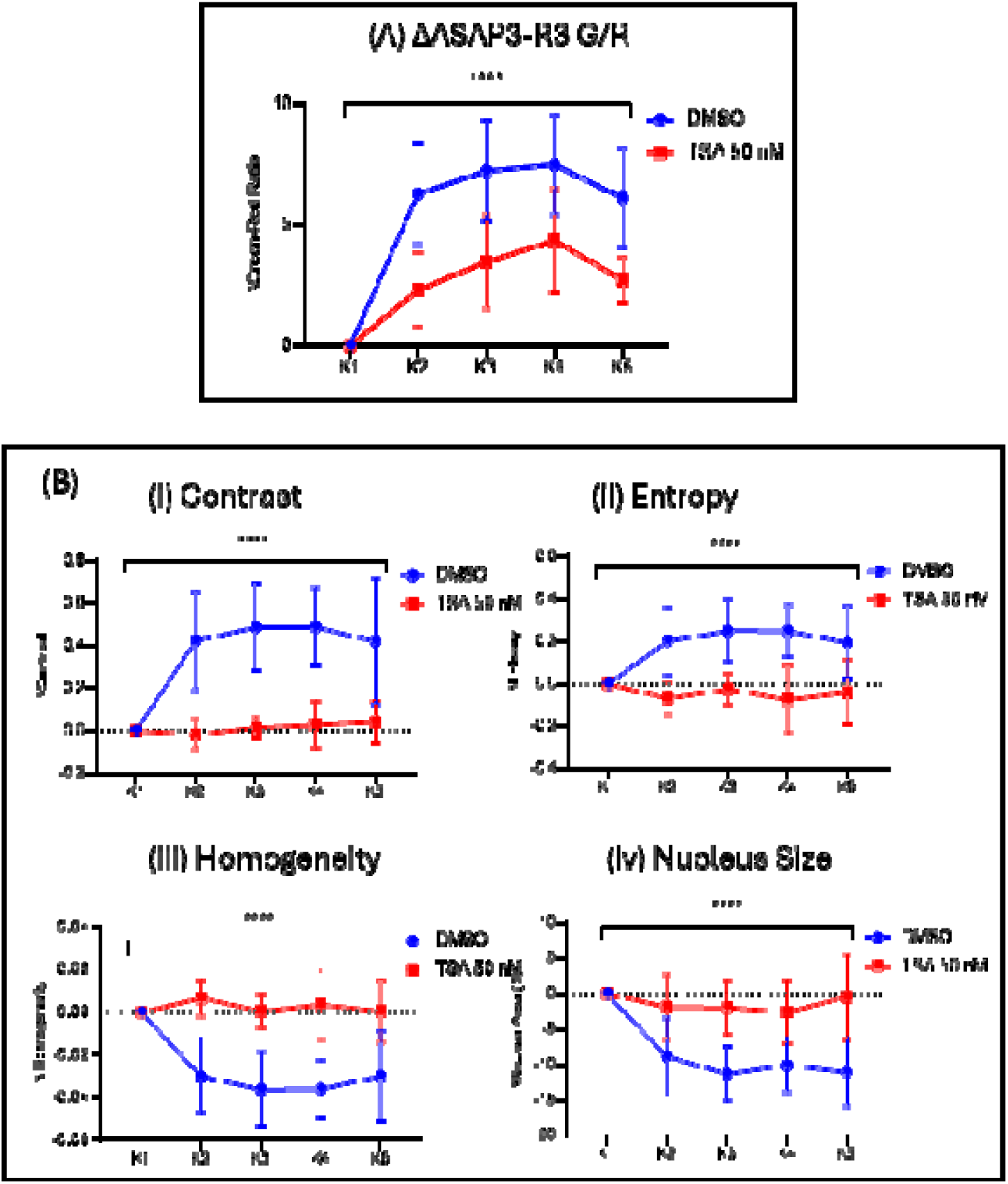
Global chromatin relaxation blunts potassium-induced V_nuc_ responses and associated chromatin dynamics. Intact NRK cells were cultured in Cellvis 96-well plates for 48 h before imaging at 37 °C and 5% CO_₂_. Cells were treated with TSA (100 nM) or an equivalent volume of DMSO for 24 h before imaging. Cells were then stained with Hoechst 33342 (1 µM) in complete FluoroBrite medium, and the medium was replaced with potassium-containing solutions. During ramp-up experiments, cells were sequentially exposed to solutions containing progressively higher potassium concentrations. (A) ASAP3-R3 measurements at the inner nuclear membrane (INM) showed that TSA treatment significantly blunted the V_nuc_ response to increasing potassium concentration relative to DMSO-treated controls (p<0.0001, n=10, two-way ANOVA). (B) GLCM analysis similarly revealed significantly blunted responses in (i) contrast, (ii) entropy, and (iii) homogeneity in TSA-treated cells (p<0.0001, n=5, two-way ANOVA). These findings are consistent with reduced chromatin compaction dynamics and diminished potassium-associated changes in chromatin organization. Error bars represent standard deviation.

### 3.9 Chromatin Compaction Blunts Potassium-Associated INM Voltage Responses

To determine whether this effect was specific to chromatin relaxation or instead reflected a broader requirement for an appropriate chromatin starting state, we next induced global chromatin compaction using sodium azide and 2-deoxy-D-glucose (2-DG). Under potassium ramping conditions, sodium azide/2-DG treatment significantly reduced the V_nuc_ response relative to control cells (p<0.0001, n=5, two-way ANOVA; Figure 10A). This treatment also significantly blunted potassium-associated changes in contrast, entropy, and homogeneity (p<0.0001, n=10, two-way ANOVA; Figure 10B-i-iii). Nuclear area analysis showed significant blunting in sodium azide/2-DG-treated cells compared with controls (p<0.0001, n=10, two-way ANOVA; Figure 10B). Together, these findings indicate that both chromatin relaxation and chromatin compaction reduce potassium-associated V_nuc_ responses and chromatin-texture dynamics.

**Figure 10.**
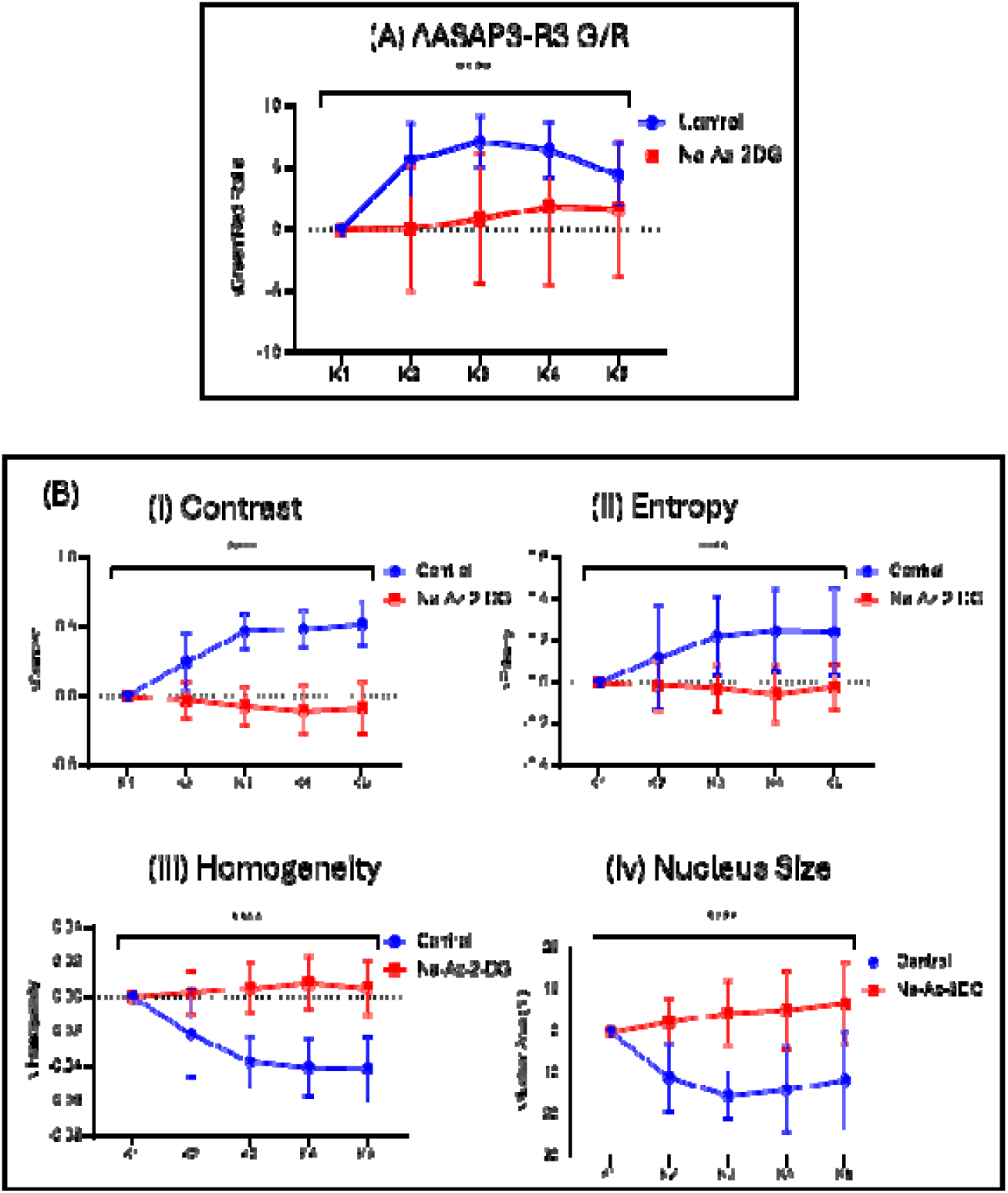
Global chromatin compaction blunts potassium-induced V_nuc_ responses and associated chromatin dynamics. Intact NRK cells were cultured in Cellvis 96-well plates for 24 h before imaging at 37 °C and 5% CO_₂_. Cells were incubated for 20 min in control complete Fluorobrite medium containing Hoechst 33342 (1 µM) alone r in incomplete DMEM media without glucose containing Hoechst 33342 (1 µM), sodium azide (10 mM), and 2-deoxy-D-glucose (2-DG; 50 mM). Cells were subsequently exposed to a ramping potassium series lacking glucose and containing sodium azide and 2-DG for an additional 10 min before imaging, whereas control cells were exposed to a glucose-containing ramping potassium series. (A) ASAP3-R3 measurements at the inner nuclear membrane (INM) showed that sodium azide^+^2-DG treatment significantly reduced the Vnuc response to increasing potassium concentrations relative to controls (p<0.0001, n=5, two-way ANOVA). (B) GLCM analysis similarly revealed significantly blunted responses in (i) contrast, (ii) entropy, and (iii) homogeneity in treated cells (p<0.0001, n=5, two-way ANOVA). Nuclear area of Na-Az ^+^ 2DG cells also showed significant blunting compared to control cells. These results are consistent with reduced potassium-associated changes in chromatin organization following induction of global chromatin compaction. Error bars represent standard deviation.

### 3.10 Chromatin Relaxation Selectively Alters Chloride-Associated INM and Chromatin Texture Responses

We next examined whether chromatin relaxation similarly altered chloride-associated responses. TSA treatment significantly reduced the chloride-induced V_nuc_ response relative to DMSO-treated controls (p<0.0001, n=5, two-way ANOVA; Figure 11A). However, blunting of chloride-induced V_nuc_ response was not significant at 50 nM like the sodium and potassium series (Supplementary Figure 1). It instead became apparent after treatment with TSA in the range of 100 - 200 nM. In the accompanying chromatin analysis, TSA significantly blunted the contrast response (p<0.0001, n=5, two-way ANOVA; Figure 11B-i), whereas entropy and homogeneity did not differ significantly between TSA-treated and control cells (Figure 11B-ii,iii). Change in nuclear area was also significantly blunted. These data indicate that chromatin relaxation suppresses chloride-evoked nuclear electrical responses, although its effects on chloride-associated chromatin-texture features are more selective than those observed for sodium or potassium.

**Figure 11.**
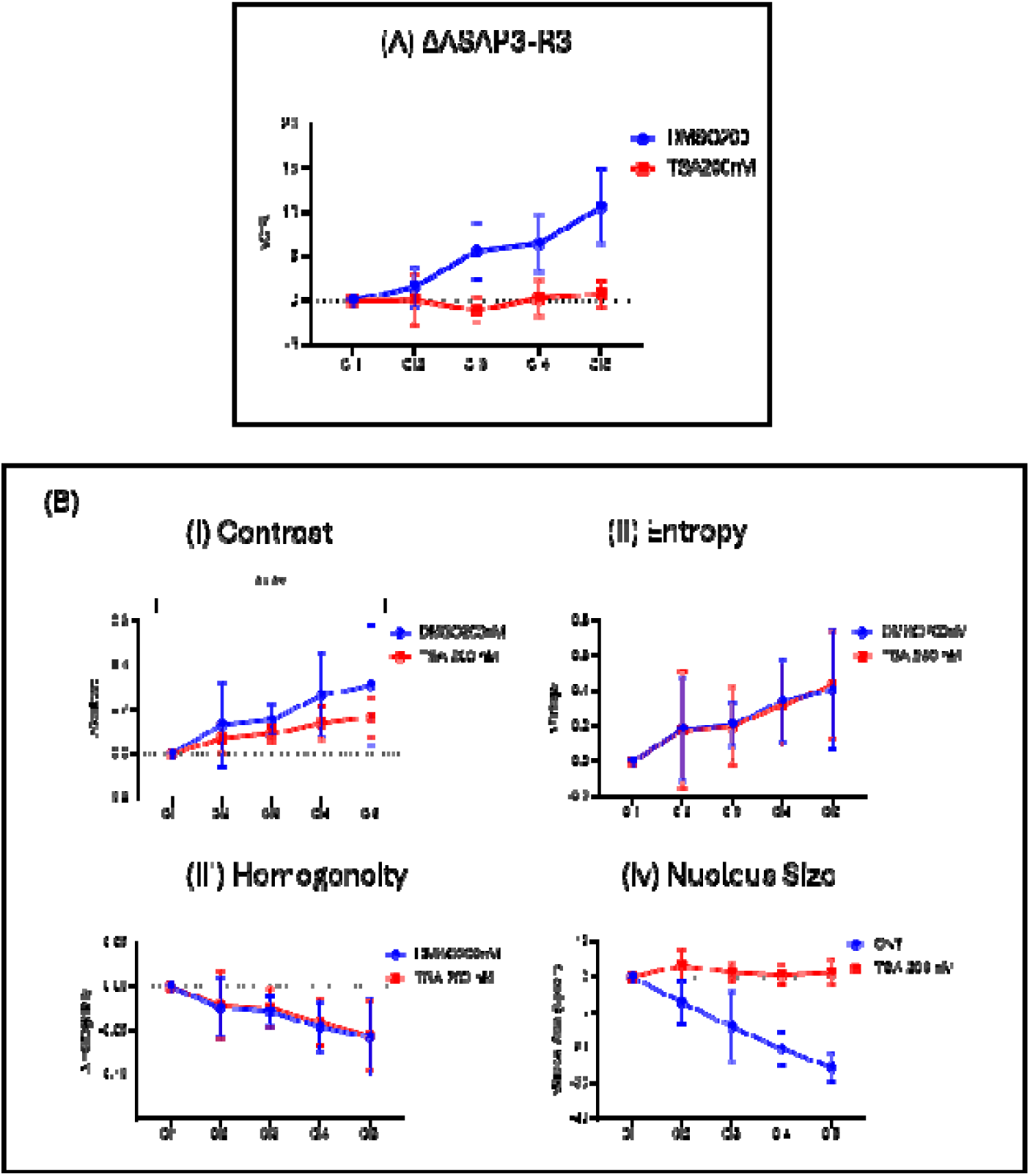
Global Relaxation blunts chloride-induced V_nuc_ responses and associated chromatin compaction. Intact NRK cells were cultured in Cellvis 96-well plates for 48 h before imaging at 37 °C and 5% CO_₂_. Cells were treated with TSA (200 nM) or an equivalent volume of DMSO for 24 h before imaging. Cells were then stained with Hoechst 33342 (1 µM) in complete FluoroBrite medium, which was directly replaced with a defined chloride solution.(A) ASAP3-R3 measurements at the inner nuclear membrane (INM) showed that TSA treatment significantly red ced the Vnuc response relative to DMSO-treated controls (p<0.0001, n=5, two-way ANOVA). (B) GLCM analysis similarly revealed a significantly blunted response in (i) contrast (p<0.0001, n=5, two-way ANOVA), whereas no significant differences were observed in (ii) entropy or (iii) homogeneity in TSA-treated cells. Together, these findings are consistent with reduced chloride-associated chromatin compaction following induction of global chromatin relaxation. Error bars represent standard deviation.

### 3.11 Chromatin Compaction Reduces Chloride-Associated INM Voltage and Chromatin Texture Remodeling

Finally, we tested whether chromatin compaction also altered chloride-associated responses. Sodium azide/2-DG treatment significantly reduced the V_nuc_ response to chloride modification relative to controls (p<0.0001, n=20, two-way ANOVA; Figure 12A). GLCM analysis revealed significantly blunted responses in contrast, entropy, and homogeneity in sodium azide/2-DG-treated cells compared with controls (p<0.0001, n=20, two-way ANOVA; Figure 12B-i-iii). Change in nuclear area was also significantly blunted in sodium azide/2-DG-treated cells compared with controls (p<0.0001, n=20, two-way ANOVA; Figure 12B). These findings show that induction of global chromatin compaction reduces chloride-associated changes in both V_nuc_ and chromatin organization.

**Figure 12.**
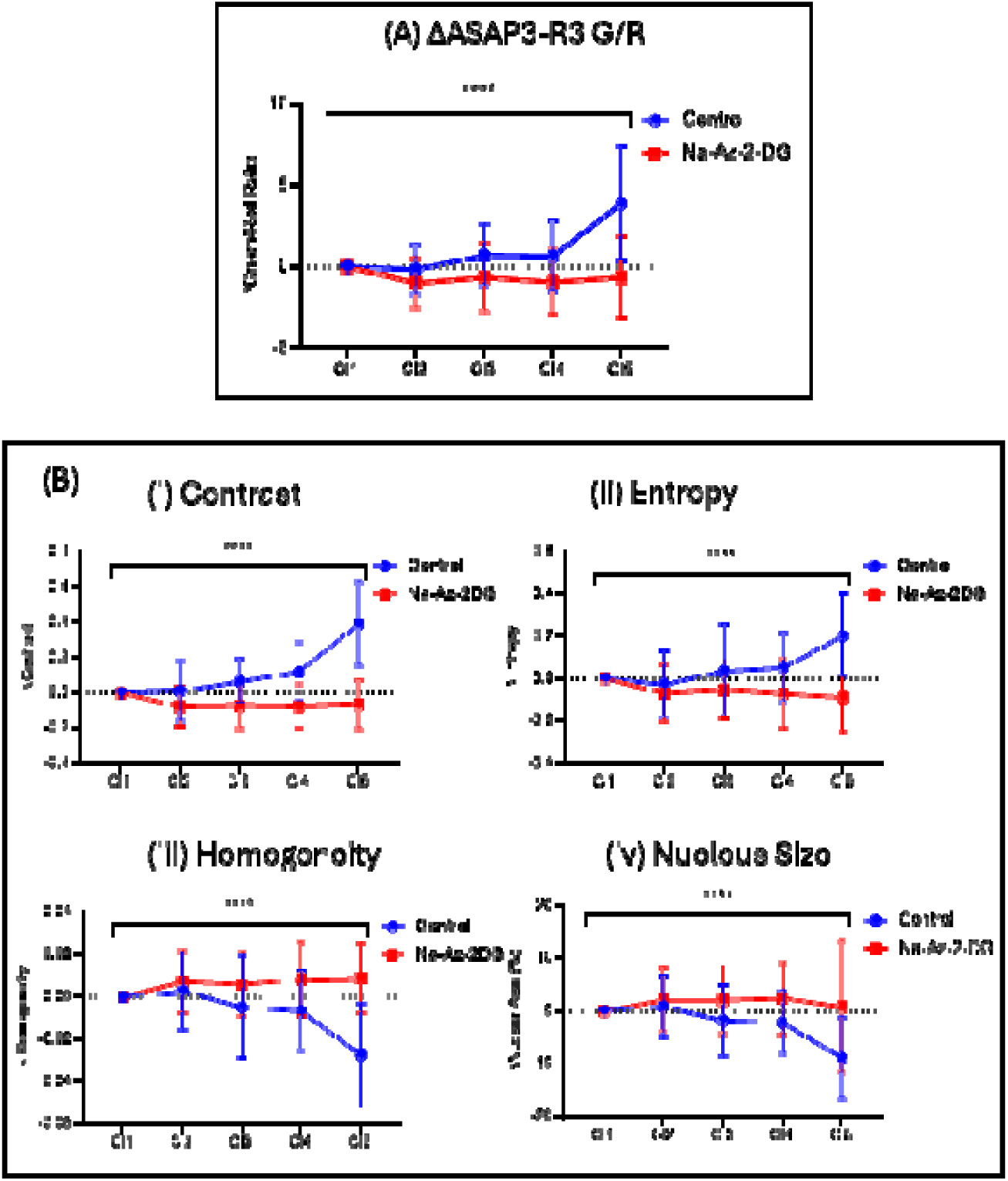
Global chromatin compaction blunts chloride-induced V_nuc_ responses and associated chromatin dynamics. Intact NRK cells were cultured in Cellvis 96-well plates for 24 h before imaging at 37 °C and 5% CO_₂_. Cells were incubated for 20 min in control complete Fluorobrite medium containing Hoechst 33342 (1 µM) alone or in incomplete DMEM media without glucose containing Hoechst 33342 (1 µM), sodium azide (10 mM), and 2-deoxy-D-glucose (2-DG; 50 mM). Cells were subsequently exposed to single chloride solution lacking glucose and containing sodium azide and 2-DG for an additional 10 min before imaging, whereas control cells were exposed to a glucose-containing chloride series. (A) ASAP3-R3 measurements at the inner nuclear membrane (INM) showed that sodiu azide^+^2-DG treatment significantly reduced the Vnuc response to increasing potassium concentrations relative to controls (p<0.0001, n=5, two-way ANOVA). (B) GLCM analysis similarly revealed significantly blunted responses in (i) contrast, (ii) entropy, and (iii) homogeneity in treated cells (p<0.0001, n=20, two-way ANOVA). Nuclear area of Na-Az ^+^ 2DG cells also showed significant blunting compared to control cells (p<0.0001, n=20, two-way ANOVA). These results are consistent with reduced chloride-associated changes in chromatin organization following induction of global chromatin compaction. Error bars represent standard deviation.

Taken together, these results define several key features of INM electro-structural responsiveness in intact NRK cells. First, the response was ion-and paradigm-dependent: sodium reduction, potassium elevation, and chloride reduction each produced INM hyperpolarization under specific conditions, but the magnitude and pattern of response depended on how the ionic perturbation was delivered (Figures 3-6). Second, potassium and sodium responses were most clearly revealed during ramping exposure, whereas chloride reduction produced responses under both direct and ramping paradigms, although the ramping response was strongest at the 30 mM chloride condition (Figures 4-6). Third, in ion conditions that produced measurable electrical responses, changes in V_nuc_ were generally accompanied by coordinated shifts in GLCM-derived chromatin features and nuclear area (Figures 3-6). Finally, experimentally shifting chromatin toward either a relaxed or compacted state blunted both the voltage response and the structural readouts across sodium, potassium, and chloride perturbations (Figures 7-12). Together, these findings suggest that INM voltage responses are both ion-dependent and chromatin-state-dependent, and they position chromatin not simply as a downstream readout, but as a major determinant of nuclear electrical responsiveness.

## Discussion

Here, we characterized how extracellular ionic perturbations influence voltage at the inner nuclear membrane (INM) and how these electrical responses relate to chromatin organization in intact NRK cells. By targeting ASAP3-R3 to SUN2-associated nuclear membranes, confirming localization by fluorescence imaging and electron microscopy (Figure 1), and functionally validating the reporter using sodium-potassium pump inhibition in isolated nuclei (Figure 2), we established a system for monitoring INM-associated voltage responses. Overall, these findings suggest that INM voltage and chromatin architecture are functionally coupled, and that this coupling depends on ionic context, exposure history, and the pre-existing chromatin state.

The localization data reveals our reporter to be situated at a membrane system that is not simply an extension of the endoplasmic reticulum (ER), but a specialized regulatory interface (Figure 1). Although the nuclear envelope is continuous with the ER, the INM has a distinct protein composition and is tightly linked to the lamina, chromatin tethering, and genome organization ^24,25,45,46^. Broad lamina-associated contacts contribute to chromosome positioning and transcriptional regulation, reinforcing the idea that membrane-localized perturbations at the nuclear periphery can be coupled to chromatin state ^25,45^. The observation that ASAP3-R3-Nuc localizes not only to the nuclear envelope but also to type I and type II nucleoplasmic reticulum invaginations is therefore notable, because the nucleoplasmic reticulum extends nuclear membrane-associated processes deeper into the nucleus and can generate local microenvironments for signaling and transport ^32^ (Figure 1). In parallel, classic electrophysiological studies established that the nuclear envelope can support ion-selective conductance and non-trivial electrical behavior, and later work identified functional Kv10.1 channels at the INM, supporting the view that voltage across this membrane system is biologically meaningful rather than merely incidental ^12–14,47^.

The isolated nuclei experiment functionally validate INM Sun2 ASAP3-R3 as a reporter of nuclear membrane potential (Figure 2). Inhibition of the INM sodium-potassium pump with strophanthidin and ouabain produced INM depolarization relative to DMSO controls, consistent with the GHK-based estimate. This supports the interpretation that the probe detects biologically meaningful INM-associated voltage changes rather than nonspecific fluorescence effects. Given that Na,K-ATPase activity has been reported at the nuclear envelope and implicated in maintaining nuclear Na^+^ and K^+^ gradients ^1^ these experiments provide the functional basis for interpreting subsequent intact-cell ionic perturbations as changes in V_Nuc_.

A notable result of this study is that the nuclear response is ion-selective, but not in a simple binary sense. The mixed sodium/potassium calibration series produced clear INM hyperpolarization and chromatin texture changes during ramping exposure, whereas direct exposure to a single endpoint condition did not significantly alter V_nuc_ or GLCM-derived chromatin texture (Figure 3). However, direct exposure did increase nuclear area, indicating that the endpoint condition was not biologically inert, even though it did not produce the same electrical or chromatin-texture response (Figure 3). Thus, the nuclear response was not determined simply by the final ionic composition. Instead, progressive ionic transitions appeared to reveal a distinct electro-structural response that was not captured by abrupt exposure.

This ordering effect is important because it suggests that V_nuc_ behaves as a dynamic state variable, dependent on a past history of physiological experiences, rather than as a simple instantaneous readout of the bathing ionic solution. In plasma membrane systems, related forms of history-dependent voltage behavior have been described as bistability, hysteresis, and bioelectric memory ^48–51^. For example, skeletal muscle fibers exposed to low extracellular potassium can occupy different membrane-potential states depending on prior ionic conditions, and Vmem can show hysteresis when extracellular potassium is gradually decreased and then increased ^48,50^. Similar K^+^-dependent bistable resting potentials have also been described in cardiac Purkinje fibers and modeled as consequences of nonlinear interactions between Na^+^/K^+^-ATPase activity, sodium conductance, and potassium conductance, particularly inwardly rectifying K+ channels ^49,52^. More broadly, bioelectric memory models propose that all cells, not only neurons, can maintain distinct voltage states depending on channel composition and prior state history ^51,53^. Although these studies concern the plasma membrane rather than the nuclear envelope, they provide an important conceptual precedent: membrane voltage can depend on the trajectory of stimulation, not only on the final ionic condition. This is likely to have significant implications for the design of intervention strategies that take advantage of the bioelectric interface to cellular and subcellular functions in biomedical contexts.

The sodium-only series helps clarify this pattern (Figure 4). Direct sodium exposure produced only limited electrical and structural effects, with V_nuc_ remaining largely unchanged except at the lowest sodium condition and with no significant changes in contrast, entropy, or homogeneity. By contrast, progressively lowering sodium during ramping produced significant INM hyperpolarization accompanied by increased contrast and entropy and reduced homogeneity, consistent with a chromatin compaction-associated texture response (Figure 4). Thus, sodium is not inert in this system, but its effect is strongly dependent on the mode of perturbation. In this context, the mixed sodium/potassium calibration ramp is best interpreted not as a pure potassium effect, but as a combined response in which stepwise sodium reduction and potassium elevation act together to move the nucleus into a distinct electro-structural state (Figures 3 and 4). By contrast, in the direct mixed condition, the abrupt endpoint condition may be insufficient to engage the same transition, or may produce competing electrical, osmotic, and chromatin effects that flatten the net V_nuc_ response. This remains an inference from the present data, rather than a directly demonstrated mechanism.

The sodium data are particularly interesting when viewed alongside chromatin biophysics (Figure 4). In reconstituted nucleosome arrays under mixed monovalent/divalent salt conditions, sodium and potassium exert different, and in some cases opposing, effects on chromatin folding, demonstrating that monovalent ions cannot be treated as interchangeable regulators of chromatin structure ^34^. Notably, Allahverdi et al. found that sodium promoted nucleosome array folding under mixed salt conditions, whereas potassium inhibited complete folding and could partially open compacted arrays ^34^. This direction differs from the present live-cell results, where increased sodium depolarized the INM and was associated with chromatin relaxation, while increased potassium hyperpolarized the INM and was associated with chromatin condensation. This divergence suggests that the chromatin responses observed here are unlikely to reflect direct cation–nucleosome interactions alone. Instead, they may emerge from the combined effects of ionic composition, INM voltage state, nuclear envelope ion handling, chromatin–lamina interactions, and the broader regulatory context of intact cells. Nucleosomes also retain a strong negative electrostatic field despite histone association, meaning that counterion interactions remain central to chromatin organization ^35^. Although these in vitro findings do not map directly onto our live-cell readouts, they provide a plausible framework for why sodium withdrawal and potassium elevation need not behave as simple mirror-image perturbations, and why sodium effects could be non-additive, non-equilibrium, and history-dependent.

A further point of relevance is that Na^+^/K^+^ gradients can exist across the nuclear envelope itself: Garner reported that Na,K-ATPase activity at the inner nuclear membrane contributes to Na^+^ and K^+^ gradients and that nuclear pores are not freely permeable to these ions ^1^. This raises the possibility that sodium perturbation may influence nuclear responses not only through direct electrostatic effects on chromatin, but also through transporter-dependent regulation of the nuclear ionic milieu and INM voltage. Functional Na^+^/K^+^-ATPase and Na^+^/Ca2^+^ exchange have also been reported at the nuclear envelope, where they contribute to nucleoplasmic Ca2^+^ homeostasis ^54^Although the present study does not test whether Na^+^/Ca2^+^ exchange, including possible reverse-mode behavior, contributes to the sodium-ramping effects observed here, such a mechanism could help explain why progressive sodium reduction produces a stronger response than abrupt sodium exposure. Together, these studies support the idea that progressive sodium change may allow the envelope-chromatin system to traverse intermediate electrochemical states, alter transporter driving forces, or engage transport-dependent compensation that abrupt exposure bypasses. Therefore, the analogy to plasma membrane bistability should be treated as functional or phenomenological rather than mechanistic at this stage. Future experiments comparing ramp-up, ramp-down, and repeated ionic cycles will be required to determine whether V_nuc_ exhibits true hysteresis beyond the order-dependent response revealed here by the difference between ramping and direct exposure.

An additional feature of the potassium data is the clear order-dependent effect (Figure 5). Direct exposure to a single potassium condition did not significantly change Vnuc, yet it altered chromatin texture and increased nuclear area. This partially overlaps with the mixed sodium/potassium calibration series, where direct exposure also increased nuclear area despite no significant change in V_nuc_ or GLCM metrics (Figure 3). Together, these direct-exposure results suggest that abrupt ionic perturbation can still influence nuclear size, even when it does not engage the full electro-structural response. In contrast, gradual ramping through increasing potassium concentrations produced robust INM hyperpolarization together with increased contrast and entropy, reduced homogeneity, and decreased nuclear area (Figure 5). Thus, the potassium response was not determined simply by the final potassium concentration, but by the sequence through which that concentration was reached. These data suggest that the nucleus is path-dependent with respect to ionic stimulation, and that progressive potassium elevation is required to engage the full electrical and chromatin compaction-associated response. One possible explanation is that sequential exposure allows the system to traverse intermediate electrochemical states that are bypassed during abrupt exposure. Another is that progressive potassium elevation more effectively recruits voltage-sensitive or state-dependent conductance at the nuclear envelope. This idea is plausible given prior reports of ion channel activity in the nuclear envelope and the localization of functional Kv10.1 channels to the INM ^13,14,47^. However, the present data do not identify the responsible channel or transporter, and that mechanistic question remains open.

The chloride data add a third important layer to the interpretation (Figure 6). In contrast to potassium and sodium, decreasing chloride produced significant graded INM hyperpolarization under direct exposure, and it also produced coordinated increases in contrast and entropy, reduced homogeneity, and decreased nuclear area. During ramping, chloride produced a more selective electrical response, with significant hyperpolarization detected at the 30 mM chloride condition, whereas the higher chloride concentrations showed a more blunted response. Chromatin texture changes during chloride ramping were also more selective, with entropy and homogeneity changing but contrast remaining unchanged (Figure 6). This suggests that chloride perturbation accesses the nuclear electrophysiological system through a response profile distinct from both sodium and potassium. That interpretation is consistent with previous reports of chloride-selective conductance in the nuclear membrane and with work identifying NCC27/CLIC-related nuclear chloride channel activity linked to cell-cycle regulation ^15,55^. More broadly, chloride is increasingly recognized as a major intracellular signaling ion involved in charge compensation, cell-volume control, pH regulation, and organelle function, including within the nucleus and ER ^2^. Accordingly, the chloride series may be acting through both a direct electrical route and a broader physicochemical route involving counterion balance, nuclear hydration, and intracellular ionic homeostasis.

The chromatin results are also notable. Across the responsive ramping conditions, INM hyperpolarization was generally associated with increased GLCM contrast and entropy together with reduced homogeneity and reduced nuclear area (Figures 3-6). While GLCM analysis does not directly resolve nucleosome packing or higher-order genome topology, the internal consistency of this signature across sodium-ramping, potassium-ramping, and chloride conditions supports the interpretation that the nucleus is shifting toward a more heterogeneous and structurally partitioned chromatin state (Figures 4-6). In practical terms, this is consistent with increased architectural heterogeneity and reduced uniformity of nuclear organization, although the exact ultrastructural substrate remains to be determined. This interpretation fits with current views of the nuclear envelope and lamina as regulators of genome architecture and with biophysical work showing that ionic conditions strongly influence chromatin folding and self-association ^25,33–35,45^. Thus, our data suggest that ion-dependent changes in V_nuc_ are accompanied by ion-dependent changes in chromatin organization, rather than representing an isolated membrane phenomenon.

Our results suggest that chromatin does not behave simply as a downstream readout of nuclear voltage. Instead, baseline chromatin state appears to shape the magnitude of the electrical response itself (Figures 7-12). TSA-induced chromatin relaxation blunted the sodium-associated V_nuc_ response, although this effect was most apparent at the two lowest sodium concentrations (Figure 7). In control cells, the sodium-ramping V_nuc_ response appeared to plateau after the 84 mM solution, including 56 mM and 28 mM, whereas TSA-treated cells showed a significantly lower response at 28 mM. This suggests that chromatin relaxation may particularly restrict the ability of the nucleus to sustain or reach the full hyperpolarizing response during stronger sodium reduction. TSA also significantly blunted potassium-associated V_nuc_ responses and reduced associated chromatin texture changes (Figure 9) ^56,57^. TSA further suppressed chloride-evoked V_nuc_ responses, although this effect was not significant at 50 nM and became apparent at higher TSA concentrations in the range of 100-200 nM (Figure 11; Supplementary Figure 1). This suggests that chloride-associated responses may require a stronger degree of chromatin relaxation before the electrical response is detectably altered, or that chloride engages additional mechanisms that are less sensitive to mild chromatin decondensation. Conversely, sodium azide plus 2-DG, which promotes ATP depletion-associated large-scale chromatin hypercompaction through altered divalent-cation and polyamine availability, blunted sodium-, potassium-, and chloride-associated responses (Figures 8, 10 and 12) ^56,58^. The fact that both chromatin relaxation and chromatin compaction reduced the magnitude of the ionic response argues against a simple one-directional model in which ions drive chromatin and chromatin merely follows. Instead, these data suggest that chromatin state sets the gain of the nuclear electrical response.

We therefore favor an operating-window model for nuclear electro-structural coupling (Figures 7-12). In this model, the nucleus is most responsive when chromatin resides within a permissive intermediate state. If chromatin is pre-relaxed, the system may lose some of the structural constraints or tethering relationships needed to convert ionic perturbation into a robust electrical and architectural transition. If chromatin is pre-compacted, the system may instead become mechanically or electrostatically constrained, leaving less dynamic range for further ion-driven reorganization. Although these possibilities remain to be tested directly, the convergence of opposite chromatin-state manipulations onto the same functional phenotype, a blunted V_nuc_ response and reduced chromatin dynamics, strongly suggests that chromatin is part of the machinery that determines nuclear responsiveness, not merely the output. This view is consistent with broader models in which the nuclear envelope, lamina, and chromatin form an integrated regulatory system rather than separable modules ^24,25,45^.

These findings also have broader implications for how the nucleus senses and interprets its ionic environment. Previous work has often treated nuclear electrophysiology and chromatin biology as related but still largely separate problems. Our data suggest that these two layers are tightly linked. Progressive sodium reduction, potassium elevation, and chloride reduction do not simply alter membrane voltage; they also produce structured changes in chromatin organization (Figures 3-6). Conversely, experimentally altering chromatin state changes how strongly the nucleus responds electrically to ionic perturbation (Figures 7-12). This supports a bidirectional model in which ionic conditions influence chromatin architecture, while chromatin architecture reciprocally constrains the amplitude of V_nuc_ responses. In this framework, V_nuc_ is not just a membrane readout, but a low-dimensional integrator of nuclear state.

### Limitations

There are several limitations that could be addressed in future work. First, the present study does not identify the channels or transport pathways responsible for the observed sodium-, potassium-, and chloride-dependent responses (Figures 3-6). Candidate-based perturbation of nuclear potassium and chloride conductance, as well as transport systems that shape nuclear Na^+^/K^+^ gradients, will therefore be important. Second, although the chromatin texture data are internally consistent, GLCM metrics remain indirect texture descriptors rather than direct measurements of chromatin folding state (Figures 3-12) ^36–38^. Orthogonal approaches such as higher-resolution structural imaging, chromatin contact assays, or direct measurements of lamina association will be needed to determine whether the observed changes reflect altered nucleosome density, domain segregation, phase behavior, or chromatin-envelope contacts. Third, the chloride experiments may reflect not only voltage effects, but also chloride-sensitive changes in pH, osmotic balance, and nuclear hydration (Figure 6) ^2^. Fourth, because the outer nuclear membrane is continuous with the ER, additional work will be needed to distinguish which component of the envelope system contributes most directly to the measured voltage signal ^24,46^. Finally, the present study does not directly test whether the changes in V_nuc_ cause chromatin reorganization, whether chromatin reorganization causes changes in V_nuc_, or whether both arise from a coupled electrochemical process. Future experiments combining voltage manipulation, ion-channel inhibition, and direct chromatin-state measurements will be needed to resolve this causal relationship.

## Conclusion

Despite these limitations, the present study establishes several key points. First, Sun2 ASAP3-R3 is both correctly localized and functionally responsive to perturbation of the INM sodium-potassium pump, supporting its use as a reporter of nuclear membrane potential (Figures 1 and 2). Second, progressive sodium reduction, potassium elevation, and chloride reduction are effective regulators of INM voltage and chromatin organization in intact cells (Figures 3-6). Third, these responses are strongly paradigm-dependent, with sodium and potassium responses emerging most clearly during ramping exposure, whereas chloride reduction produces a more direct response profile (Figures 4-6). Fourth, ion conditions that induce INM hyperpolarization are generally associated with increased chromatin contrast and entropy, reduced homogeneity, and reduced nuclear area, consistent with increased chromatin architectural complexity and compaction-associated reorganization (Figures 3-6). Finally, both chromatin relaxation and chromatin compaction blunt ion-evoked V_nuc_ responses, indicating that baseline chromatin state gates nuclear electrical responsiveness (Figures 7-12). Taken together, these data support a model in which the nuclear envelope and chromatin form a coupled electro-structural system that interprets ionic conditions and translates them into changes in nuclear organization. More broadly, they suggest that membrane voltage at the nuclear envelope may serve as an important biophysical variable linking ion homeostasis, chromatin architecture, and nuclear function.

## Acknowledgements

We thank Patrick McMillen for the confocal training he provided. This work was funded partially by the Australian Fulbright Commission, a Sponsored Research Agreement to Tufts University from Astonishing Labs, and by grant 62212 from the John Templeton Foundation. The opinions expressed in this publication are those of the author(s) and do not necessarily reflect the views of the John Templeton Foundation nor Fulbright. This work was also supported in part by the Koch Institute Support (core) Grant P30-CA14051 from the National Cancer Institute. We thank the Koch Institute’s Robert A. Swanson (1969) Biotechnology Center for support, specifically Peterson (1957) Nanotechnology Materials Core Facility (RRID:SCR_018674).

---

**Supplementary Figure 1.**
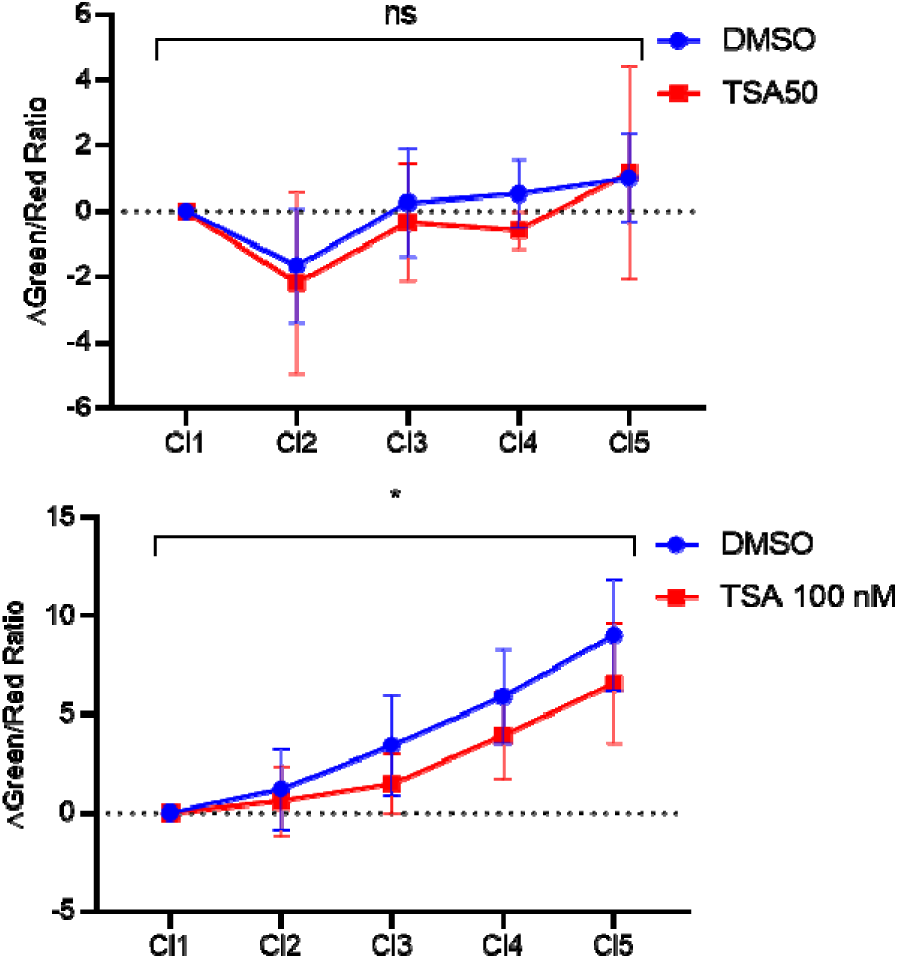
TSA concentrations optimization for blunting Chloride response. Intact NRK cells were cultured in Cellvis 96-well plates for 48 h before imaging at 37 °C and 5% CO_₂_. Cells were treated with TSA (200 nM) or an equivalent volume of DMSO for 24 h before imaging. Cells were then stained with Hoechst 33342 (1 µM) in complete FluoroBrite medium, which was directly replaced with a defined chloride solution. (A) ASAP3-R3 measurements at the inner nuclear membrane (INM) showed that TSA treatment (50 nM) did not significantly blunt the Vnuc response relative to DMSO-treated controls (p>0.05, n=5, two-way ANOVA). However, significant blunting Vnuc response to chloride series was observed with 100 nM TSA treatment. Error bars represent standard deviation.

## References

1. Garner, M. (2002). Na, K-ATPase in the nuclear envelope regulates Na+: K+ gradients in hepatocyte nuclei. J. Membr. Biol. 187, 97–115.

2. Satish, Singh, K., Sanghvi, S., Loyo-Celis, V., Varghese, L., Ekam, Gururaja Rao, S., and Singh, H. (2024). Chloride ions in health and disease. Biosci. Rep. 44. 10.1042/bsr20240029.

3. George, L.F., and Bates, E.A. (2022). Mechanisms Underlying Influence of Bioelectricity in Development. Front Cell Dev Biol 10, 772230. 10.3389/fcell.2022.772230.

4. Levin, M. (2021). Bioelectric signaling: Reprogrammable circuits underlying embryogenesis, regeneration, and cancer. Cell 184, 1971–1989. 10.1016/j.cell.2021.02.034.

5. McLaughlin, K.A., and Levin, M. (2018). Bioelectric signaling in regeneration: Mechanisms of ionic controls of growth and form. Dev. Biol. 433, 177–189. 10.1016/j.ydbio.2017.08.032.

6. Silic, M.R., and Zhang, G. (2023). Bioelectricity in Developmental Patterning and Size Control: Evidence and Genetically Encoded Tools in the Zebrafish Model. Cells 12. 10.3390/cells12081148.

7. Harris, M.P. (2021). Bioelectric signaling as a unique regulator of development and regeneration. Development 148. 10.1242/dev.180794.

8. Chernet, B.T., and Levin, M. (2013). Transmembrane voltage potential is an essential cellular parameter for the detection and control of tumor development in a Xenopus model. Dis. Model. Mech. 6, 595–607. 10.1242/dmm.010835.

9. Levin, M., Pezzulo, G., and Finkelstein, J.M. (2017). Endogenous Bioelectric Signaling Networks: Exploiting Voltage Gradients for Control of Growth and Form. Annu. Rev. Biomed. Eng. 19, 353–387. 10.1146/annurev-bioeng-071114-040647.

10. Hodgkin, A.L., and Katz, B. (1949). The effect of sodium ions on the electrical activity of giant axon of the squid. J. Physiol. 108, 37–77. 10.1113/jphysiol.1949.sp004310.

11. Matzke, A.J., Weiger, T.M., and Matzke, M. (2010). Ion channels at the nucleus: electrophysiology meets the genome. Molecular plant 3, 642–652.

12. Loewenstein, W.R., and Kanno, Y. (1963). Some Electrical Properties of a Nuclear Membrane Examined with a Microelectrode. The Journal of General Physiology 46, 1123–1140. 10.1085/jgp.46.6.1123.

13. Mazzanti, M., Bustamante, J.O., and Oberleithner, H. (2001). Electrical dimension of the nuclear envelope. Physiol. Rev. 81, 1–19.

14. Mazzanti, M., DeFelice, L.J., Cohen, J., and Malter, H. (1990). Ion channels in the nuclear envelope. Nature 343, 764. 10.1038/343764a0.

15. Tabares, L., Mazzanti, M., and Clapham, D.E. (1991). Chloride channels in the nuclear membrane. J. Membr. Biol. 123, 49–54. 10.1007/bf01993962.

16. Rousseau, E., Michaud, C., Lefebvre, D., Proteau, S., and Decrouy, A. (1996). Reconstitution of ionic channels from inner and outer membranes of mammalian cardiac nuclei. Biophys. J. 70, 703–714. 10.1016/s0006-3495(96)79610-8.

17. Secondo, A., Esposito, A., Petrozziello, T., Boscia, F., Molinaro, P., Tedeschi, V., Pannaccione, A., Ciccone, R., Guida, N., Di Renzo, G., and Annunziato, L. (2018). Na(+)/Ca(2+) exchanger 1 on nuclear envelope controls PTEN/Akt pathway via nucleoplasmic Ca(2+) regulation during neuronal differentiation. Cell Death Discov 4, 12. 10.1038/s41420-017-0018-1.

18. Wente, S.R., and Rout, M.P. (2010). The nuclear pore complex and nuclear transport. Cold Spring Harb. Perspect. Biol. 2, a000562. 10.1101/cshperspect.a000562.

19. Frey, S., and Görlich, D. (2007). A saturated FG-repeat hydrogel can reproduce the permeability properties of nuclear pore complexes. Cell 130, 512–523. 10.1016/j.cell.2007.06.024.

20. Lowe, A.R., Tang, J.H., Yassif, J., Graf, M., Huang, W.Y., Groves, J.T., Weis, K., and Liphardt, J.T. (2015). Importin-β modulates the permeability of the nuclear pore complex in a Ran-dependent manner. eLife 4. 10.7554/eLife.04052.

21. Kim, J., Izadyar, A., Shen, M., Ishimatsu, R., and Amemiya, S. (2014). Ion permeability of the nuclear pore complex and ion-induced macromolecular permeation as studied by scanning electrochemical and fluorescence microscopy. Anal. Chem. 86, 2090–2098. 10.1021/ac403607s.

22. Pathirathna, P., Balla, R.J., Jantz, D.T., Kurapati, N., Gramm, E.R., Leonard, K.C., and Amemiya, S. (2019). Probing High Permeability of Nuclear Pore Complexes by Scanning Electrochemical Microscopy: Ca(2+) Effects on Transport Barriers. Anal. Chem. 91, 5446–5454. 10.1021/acs.analchem.9b00796.

23. Zimmerli, C.E., Allegretti, M., Rantos, V., Goetz, S.K., Obarska-Kosinska, A., Zagoriy, I., Halavatyi, A., Hummer, G., Mahamid, J., Kosinski, J., and Beck, M. (2021). Nuclear pores dilate and constrict in cellulo. Science 374, eabd9776. 10.1126/science.abd9776.

24. Hetzer, M.W. (2010). The nuclear envelope. Cold Spring Harb. Perspect. Biol. 2, a000539. 10.1101/cshperspect.a000539.

25. Guerreiro, I., and Kind, J. (2019). Spatial chromatin organization and gene regulation at the nuclear lamina. Curr Opin Genet Dev 55, 19–25. 10.1016/j.gde.2019.04.008.

26. Marshall, W.F., Straight, A., Marko, J.F., Swedlow, J., Dernburg, A., Belmont, A., Murray, A.W., Agard, D.A., and Sedat, J.W. (1997). Interphase chromosomes undergo constrained diffusional motion in living cells. Curr. Biol. 7, 930–939. 10.1016/s0960-9822(06)00412-x.

27. Chubb, J.R., Boyle, S., Perry, P., and Bickmore, W.A. (2002). Chromatin motion is constrained by association with nuclear compartments in human cells. Curr. Biol. 12, 439–445. 10.1016/s0960-9822(02)00695-4.

28. Hinde, E., Cardarelli, F., Digman, M.A., and Gratton, E. (2010). In vivo pair correlation analysis of EGFP intranuclear diffusion reveals DNA-dependent molecular flow. Proc. Natl. Acad. Sci. U. S. A. 107, 16560–16565. 10.1073/pnas.1006731107.

29. Hinde, E., Cardarelli, F., Digman, M.A., and Gratton, E. (2012). Changes in chromatin compaction during the cell cycle revealed by micrometer-scale measurement of molecular flow in the nucleus. Biophys. J. 102, 691–697. 10.1016/j.bpj.2011.11.4026.

30. Rashid, F., Kabbo, S.A., and Wang, N. (2024). Mechanomemory of nucleoplasm and RNA polymerase II after chromatin stretching by a microinjected magnetic nanoparticle force. Cell Rep 43, 114462. 10.1016/j.celrep.2024.114462.

31. Sabari, B.R., Dall’Agnese, A., and Young, R.A. (2020). Biomolecular Condensates in the Nucleus. Trends Biochem. Sci. 45, 961–977. 10.1016/j.tibs.2020.06.007.

32. Malhas, A., Goulbourne, C., and Vaux, D.J. (2011). The nucleoplasmic reticulum: form and function. Trends Cell Biol. 21, 362–373. 10.1016/j.tcb.2011.03.008.

33. Korolev, N., Allahverdi, A., Yang, Y., Fan, Y., Lyubartsev, A.P., and Nordenskiöld, L. (2010). Electrostatic origin of salt-induced nucleosome array compaction. Biophys. J. 99, 1896–1905. 10.1016/j.bpj.2010.07.017.

34. Allahverdi, A., Chen, Q., Korolev, N., and Nordenskiöld, L. (2015). Chromatin compaction under mixed salt conditions: opposite effects of sodium and potassium ions on nucleosome array folding. Sci. Rep. 5, 1–7.

35. Gebala, M., Johnson, S.L., Narlikar, G.J., and Herschlag, D. (2019). Ion counting demonstrates a high electrostatic field generated by the nucleosome. eLife 8. 10.7554/eLife.44993.

36. Haralick, R.M., Shanmugam, K., and Dinstein, I.H. (1973). Textural features for image classification. IEEE Transactions on systems, man, and cybernetics, 610-621.

37. Dinčić, M., Todorović, J., Nešović Ostojić, J., Kovačević, S., Dunđerović, D., Lopičić, S., Spasić, S., Radojević-Škodrić, S., Stanisavljević, D., and Ilić, A.Ž. (2020). The Fractal and GLCM Textural Parameters of Chromatin May Be Potential Biomarkers of Papillary Thyroid Carcinoma in Hashimoto’s Thyroiditis Specimens. Microsc. Microanal. 26, 717–730. 10.1017/S1431927620001683.

38. Lee, H.K., Kim, C.H., Bhattacharjee, S., Park, H.G., Prakash, D., and Choi, H.K. (2021). A Paradigm Shift in Nuclear Chromatin Interpretation: From Qualitative Intuitive Recognition to Quantitative Texture Analysis of Breast Cancer Cell Nuclei. Cytometry A 99, 698–706. 10.1002/cyto.a.24260.

39. Kim, B.B., Wu, H., Hao, Y.A., Pan, M., Chavarha, M., Zhao, Y., Westberg, M., St-Pierre, F., Wu, J.C., and Lin, M.Z. (2022). A red fluorescent protein with improved monomericity enables ratiometric voltage imaging with ASAP3. Sci. Rep. 12, 3678. 10.1038/s41598-022-07313-1.

40. Matzke, A.J.M., and Matzke, M. (2015). Expression and testing in plants of ArcLight, a genetically–encoded voltage indicator used in neuroscience research. BMC Plant Biol. 15, 245. 10.1186/s12870-015-0633-z.

41. Hodzic, D.M., Yeater, D.B., Bengtsson, L., Otto, H., and Stahl, P.D. (2004). Sun2 is a novel mammalian inner nuclear membrane protein. J. Biol. Chem. 279, 25805–25812. 10.1074/jbc.M313157200.

42. Marh, J., Stoytcheva, Z., Urschitz, J., Sugawara, A., Yamashiro, H., Owens, J.B., Stoytchev, I., Pelczar, P., Yanagimachi, R., and Moisyadi, S. (2012). Hyperactive self-inactivating piggyBac for transposase-enhanced pronuclear microinjection transgenesis. Proc. Natl. Acad. Sci. U. S. A. 109, 19184–19189. 10.1073/pnas.1216473109.

43. Owens, J.B., Mathews, J., Davy, P., Stoytchev, I., Moisyadi, S., and Allsopp, R. (2013). Effective Targeted Gene Knockdown in Mammalian Cells Using the piggyBac Transposase-based Delivery System. Mol Ther Nucleic Acids 2, e137. 10.1038/mtna.2013.61.

44. Bonzanni, M., Payne, S.L., Adelfio, M., Kaplan, D.L., Levin, M., and Oudin, M.J. (2020). Defined extracellular ionic solutions to study and manipulate the cellular resting membrane potential. Biol Open 9. 10.1242/bio.048553.

45. Dechat, T., Pfleghaar, K., Sengupta, K., Shimi, T., Shumaker, D.K., Solimando, L., and Goldman, R.D. (2008). Nuclear lamins: major factors in the structural organization and function of the nucleus and chromatin. Genes Dev 22, 832–853. 10.1101/gad.1652708.

46. Schirmer, E.C., and Gerace, L. (2005). The nuclear membrane proteome: extending the envelope. Trends Biochem. Sci. 30, 551–558. 10.1016/j.tibs.2005.08.003.

47. Chen, Y., Sánchez, A., Rubio, M.E., Kohl, T., Pardo, L.A., and Stühmer, W. (2011). Functional KV10.1 Channels Localize to the Inner Nuclear Membrane. PLoS One 6, e19257. 10.1371/journal.pone.0019257.

48. Geukes Foppen, R.J., van Mil, H.G., and van Heukelom, J.S. (2002). Effects of chloride transport on bistable behaviour of the membrane potential in mouse skeletal muscle. J. Physiol. 542, 181–191. 10.1113/jphysiol.2001.013298.

49. van Mil, H., Siegenbeek van Heukelom, J., and Bier, M. (2003). A bistable membrane potential at low extracellular potassium concentration. Biophys. Chem. 106, 15–21. 10.1016/s0301-4622(03)00135-2.

50. Gallaher, J., Bier, M., and van Heukelom, J.S. (2010). First order phase transition and hysteresis in a cell’s maintenance of the membrane potential—An essential role for the inward potassium rectifiers. Biosystems 101, 149–155. 10.1016/j.biosystems.2010.05.007.

51. Law, R., and Levin, M. (2015). Bioelectric memory: modeling resting potential bistability in amphibian embryos and mammalian cells. Theor. Biol. Med. Model. 12, 22. 10.1186/s12976-015-0019-9.

52. Gadsby, D.C., and Cranefield, P.F. (1977). Two levels of resting potential in cardiac Purkinje fibers. The Journal of general physiology 70, 725.

53. Cervera, J., Alcaraz, A., and Mafe, S. (2014). Membrane potential bistability in nonexcitable cells as described by inward and outward voltage-gated ion channels. The Journal of Physical Chemistry B 118, 12444–12450.

54. Galva, C., Artigas, P., and Gatto, C. (2012). Nuclear Na+/K+-ATPase plays an active role in nucleoplasmic Ca2+ homeostasis. J. Cell Sci. 125, 6137–6147.

55. Valenzuela, S.M., Mazzanti, M., Tonini, R., Qiu, M.R., Warton, K., Musgrove, E.A., Campbell, T.J., and Breit, S.N. (2000). The nuclear chloride ion channel NCC27 is involved in regulation of the cell cycle. J. Physiol. 529 *Pt* *3*, 541–552. 10.1111/j.1469-7793.2000.00541.x.

56. Llères, D., James, J., Swift, S., Norman, D.G., and Lamond, A.I. (2009). Quantitative analysis of chromatin compaction in living cells using FLIM-FRET. J. Cell Biol. 187, 481–496. 10.1083/jcb.200907029.

57. Tóth, K.F., Knoch, T.A., Wachsmuth, M., Frank-Stöhr, M., Stöhr, M., Bacher, C.P., Müller, G., and Rippe, K. (2004). Trichostatin A-induced histone acetylation causes decondensation of interphase chromatin. J. Cell Sci. 117, 4277–4287.

58. Visvanathan, A., Ahmed, K., Even-Faitelson, L., Lleres, D., Bazett-Jones, D.P., and Lamond, A.I. (2013). Modulation of higher order chromatin conformation in mammalian cell nuclei can be mediated by polyamines and divalent cations. PLoS One 8, e67689.

